# Targeted mutagenesis of the *Arabidopsis* GROWTH-REGULATING FACTOR (GRF) gene family suggests competition of multiplexed sgRNAs for Cas9 apoprotein

**DOI:** 10.1101/2020.08.16.253203

**Authors:** Juan Angulo, Christopher P Astin, Olivia Bauer, Kelan J Blash, Natalee M Bowen, Nneoma J Chukwudinma, Austin S Dinofrio, Donald O Faletti, Alexa M Ghulam, Cloe M Gusinde-Duffy, Kamaria J Horace, Andrew M Ingram, Kylie E Isaack, Geon Jeong, Randolph J Kiser, Jason S Kobylanski, Madeline R Long, Grace A Manning, Julie M Morales, Kevin H Nguyen, Robin T Pham, Monthip H Phillips, Tanner W Reel, Jenny E Seo, Hiep D Vo, Alexander M Wukuson, Kathryn A Yeary, Grace Y Zheng, Wolfgang Lukowitz

**Affiliations:** University of Georgia, Department of Plant Biology, Athens, USA

**Keywords:** CRISPR, T-DNA vector, transgene-free, seed bio-fluorescence selection, LED illumination, teaching lab, citizen science, *Bsal*, sgRNA multiplexing, tRNA buffer, EC1

## Abstract

Genome editing in plants typically relies on T-DNA plasmids that are mobilized by Agrobacterium-mediated transformation to deliver the CRISPR/Cas9 machinery. Here, we introduce a series of CRISPR/Cas9 T-DNA vectors for minimal lab settings, such as in the classroom or citizen science projects. Spacer sequences targeting genes of interest can be inserted as annealed short oligonucleotides in a single straightforward cloning step. Fluorescent markers expressed in mature seeds enable reliable selection of transgenic as well as transgene-free individuals using a combination of inexpensive LED lamps and colored-glass alternative filters. Testing these tools on the *Arabidopsis* GROWTH-REGULATING FACTOR (GRF) gene family, we found that Cas9 expression from an EGG CELL1 (EC1) promoter resulted in tenfold lower mutation rates than expression from a UBIQUITIN10 (UBQ10) promoter. A collection of *bona fide* null mutations in all nine GRF genes could be established with little effort. Finally, we explored the effects of simultaneously targeting two, four and eight GRF genes on the rate of induced mutations at each target locus. Multiplexing caused strong interference effects: while mutation rates at some loci remained consistently high, mutation rates at other loci dropped dramatically with increasing number of single guide RNA species. Our results suggest potential detrimental genetic interaction between induced mutations as well as competition of CRISPR RNAs for a limiting amount of Cas9 apoprotein.

## Introduction

CRISPR/Cas complexes can be programed to bind virtually any DNA sequence and thus enable applications as diverse as chemical modification of target DNA, directed manipulation of gene transcription, or genome editing by homologous recombination (for an overview of recent work in plants see Ma & al., 2016; Soyars & al., 2018; Manghwar & al., 2019; Zhang & al., 2019). Natural CRISPR/Cas systems function predominantly as sequence-specific endo-nucleases that help bacterial cells clear invading plasmids or phages (Terns & Terns, 2011). Not surprisingly, the first and arguably one of the most valuable technical applications was to re-purpose this activity for mutagenizing specific loci in a genome of interest. The type II CRISPR-system of *Streptococcus pyogenes* can be reduced to two components: a multi-domain Cas9 apo-protein and a single guide RNA (sgRNA, a fusion of the two RNA components found in natural complexes; Jinek & al., 2012). As in bacteria, sgRNA/Cas9 complexes assembled *in vitro* or expressed in a target organism are homology-guided endo-nucleases: the 5’-most ∼20 nucleotides of the sgRNA, called spacer, are free to hybridize with complementary sequences, called protospacer, in the target genome; if the protospacer sequence is followed by a short protospacer-associated motif (PAM; the minimal Cas9 consensus is ‘NGG’), Cas9 induces a double-strand-break in the target DNA (three nucleotides upstream of the protospacer/PAM junction, leaving blunt ends). In most cell types, such lesions are repaired by non-homologous end joining, an error-prone process that frequently introduces small deletions or insertions.

We were interested in using CRISPR/Cas9 nucleases as part of a plant molecular biology teaching lab and needed to simplify the experimental workflow as much as possible. A series of T-DNA vectors that were designed for adding spacer sequences targeting genes of interest as short, synthetic oligonucleotide-assemblies in a single cloning step (Xing & al., 2014; Wang & al., 2015) provided an attractive platform for this purpose. We modified these vectors by replacing their antibiotic selection marker with makers enabling selection and counter-selection of transgenic plants on the basis of seed fluorescence (similar to Gao & al., 2014). In addition, we verified that seed fluorescence could be detected with standard dissecting microscopes using an inexpensive external illumination consisting of high-intensity LED lights and colored-glass alternative emission filters.

GROWTH-REGULATING FACTORs (GRFs) encode DNA-binding proteins that associate with GRF-INTERACTING FACTORs (GIFs) to regulate gene transcription, presumably by recruiting chromatin remodelers (reviewed in Omidbakhshfard & al., 2015; Kim, 2019). To date, GRF genes have only been found in the genomes of land plants and their algal precursors (together forming the streptophyte clade). The nine GRF genes of *Arabidopsis* are thought to affect cell division and expansion in the context of various developmental processes, but their genetic analysis is hampered by functional overlap and, in many cases, the lack of *bona fide* null alleles. Using our tools in the teaching lab, we were able to establish a collection of reference alleles truncating the predicted proteins prior to the DNA-binding domain. When we multiplexed sgRNAs to simultaneously target two, four, and eight GRF genes we observed vastly different rates of CRISPR/Cas9-induced lesions at different target loci, suggesting strong interference effects.

## Results

### CRISPR/Cas9 T-DNA vectors for selecting & counter-selecting transgenic seeds on the basis of bio-fluorescence

T-DNAs of the Cambia family (www.cambia.org; derived from pPZP; Hajdukiewicz & al., 1994) are among the most widely used plasmid vectors for plant transformation. Molecular cloning is greatly facilitated by their relatively small size and high copy numbers in *E. coli*; their pVP1 origin ensures effective propagation in *Agrobacterium* and high transformation rates with a wide range of plant species. Building on the pHEE401E plasmid, a Cambia T-DNA adapted for genome editing with CRISPR/Cas9 nucleases (Wang & al., 2015), we created a series of vectors enabling selection and counter-selection of transgenics on the basis of seed fluorescence. Toward this end, we replaced the hygromycin resistance marker of pHEE401E with a nuclear-localized fluorescent protein expressed from the promoter of the *Arabidopsis* seed storage albumin A1 gene (CRU3, At4g28520; Fig 1). Three different color variants express a cyan fluorescent protein (CFP; Cubitt & al., 1999), a ‘Venus’ yellow fluorescent protein (YFP, Nagai & al., 2002; both modified from *Aequorea*) and a monomeric ‘Tomato’ red fluorescent protein (Shaner & al., 2004; modified from *Discosoma*).

**Fig 1.**
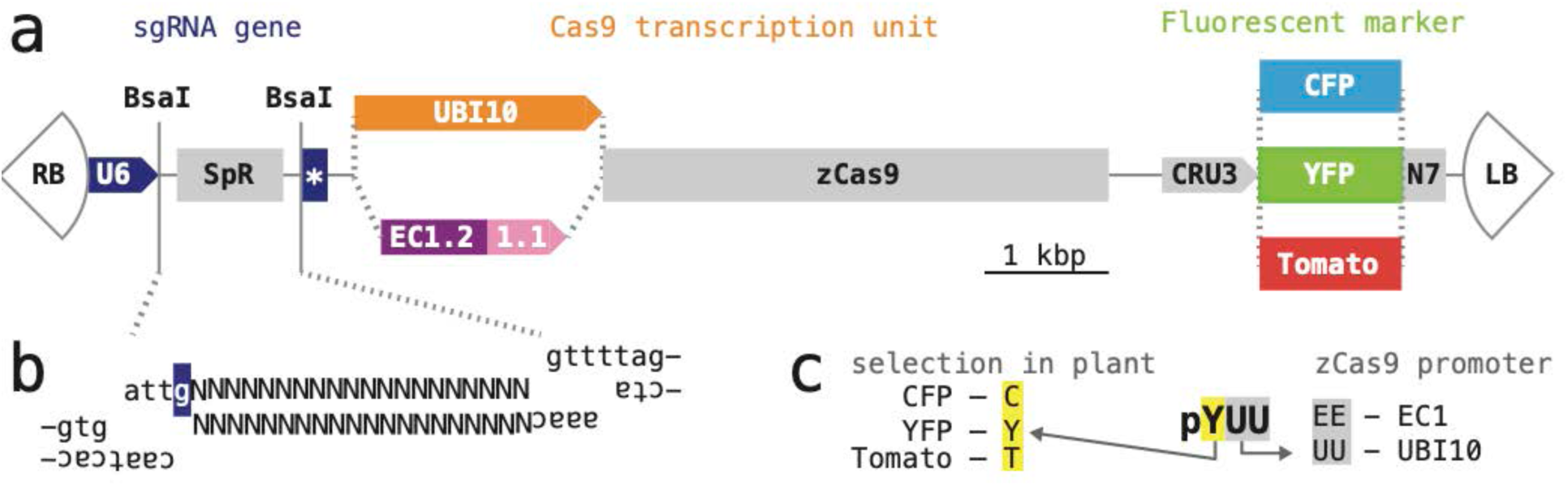
CRISPR/Cas9 T-DNAs for selection and counter-selection on the basis of seed fluorescence. (**a**) Schematic organization of T-DNA vectors. All plasmids are variants of the vectors developed by Xing & al., 2014 and Wang & al., 2015. ‘RB’ and ‘LB’, right and left T-DNA border; ‘U6’, U6-26 promoter; ‘SpR’, buffer sequence; star, sgRNA scaffold and U6 terminator; ‘UBI10’, polyubiquitin10 promoter; ‘EC1.2 1.1’, egg cell1.1 promoter with egg cell1.2 enhancer; ‘zCas9’, Cas9 coding sequence optimized for maize; ‘CRU3’, cruciferin 3 promoter; ‘YFP’, ‘CFP’, ‘Tomato’, yellow, cyan, red fluorescent protein coding sequence; ‘N7’, nuclear localization signal of At4g19150 (Cutler & al., 2000). (**b**) Annealed oligonucleotides encoding a single gene-specific spacer sequence (represented with ‘Ns’) or more complex assemblies for expression of multiple sgRNAs for the same T-DNA can be inserted into the *BsaI* cloning site; the ends generated by *BsaI* have different 5’ overhangs; the ‘g’ highlighted in dark blue represents the transcriptional start site of the U6-26 promoter. (**c**) Vectors are named after the fluorescent marker (first letter) and the promoter driving Cas9 (remaining two letters).

The CRISPR/Cas9 module of pHEE401E contains two genes, one producing the sgRNA and one producing the Cas9 mRNA. The CRISPR transcription unit consists of the *Arabidopsis* U6-26 (At3g13855) polymerase III-dependent promoter, followed by a buffer segment (SpR), a 75 bp sequence encoding the sgRNA scaffold, and the *Arabidopsis* U6-26 terminator. The buffer segment is designed to be removed by digestion with *BsaI*, a restriction enzyme cutting outside of its recognition sequence (GGTCTCN1/5). *BsaI* digestion leaves incompatible 5’-overhangs precisely at the transcriptional start site of the sgRNA gene, enabling insertion of a synthetic 19-20 base pair spacer targeting a gene of interest in a single, technically straightforward cloning step; larger fragments for expression of multiple sgRNA species may be inserted in a similar manner (Xing & al., 2014; Wang & al. 2015; Fig 1). No changes were made to this part of the T-DNA.

The Cas9 transcription unit of pHEE401E consists of egg cell-specific promoter and enhancer elements taken from the *Arabidopsis* EC1.1 and EC1.2 genes (At1g76750 and At2g21740) followed by an open reading frame optimized for maize codon usage (zCas9). Ideally, egg cell-specific activity of zCas9 would induce mutations very early in embryonic development, generating primary transgenics that are heterozygous or bi-allelic mutant rather than mosaic (Wang & al., 2015). However, pHEE401E-derived constructs tended to induce mutations at a low rate in our hands. We therefore tested an alternative promoter for zCas9, taken from the *Arabidopsis* polyubiquitin10 gene (At4g05320); this promoter drives robust transcription in a broad range of cell types and is commonly used for gene editing (Ma & al., 2016; Soyars & al., 2018; similarly, some of the plasmids described in Xiang & al., 2014, contain a maize ubiquitin1 promoter; Fig 1).

Annotated sequence listings of the six CRISPR/Cas9 T-DNA vectors can be found in the supporting material (S1 File), and samples of all plasmids have been deposited with Addgene (see Methods).

### Low-cost LED illumination for detecting bio-fluorescence in mature seeds

Ease of use is an attractive feature of bio-fluorescence markers, particularly in the context of teaching or citizen science; however, commercial fluorescence illuminations can be prohibitively expensive for such settings. Motivated by a note in ‘The Worm Breeder’s Gazette’ (Chin-Sang & Zhong, 2008), we explored the performance of high-intensity LED lights and colored-glass alternative emission filters as a means of providing external epi-fluorescence illumination for standard dissecting microscopes. We tried six LED assemblies producing relatively narrow spectra of light with maxima ranging from 415–540 nm in combination with six longpass emission filters that had cut-off values ranging from 475–610 nm (Fig 2a). Our test sample was a collection of wild type seeds spiked with a small number of seeds expressing either CFP, YFP, or Tomato from one of the T-DNAs described above (Fig 2b). As a benchmark, we imaged the same sample using an Olympus SZX12 stereo-microscope fitted with an internal fluorescence illumination module (see Methods; YFP was imaged using a GFP filter cube, Tomato using a propidium iodide filter cube; no appropriate filters were available for CFP).

**Fig 2.**
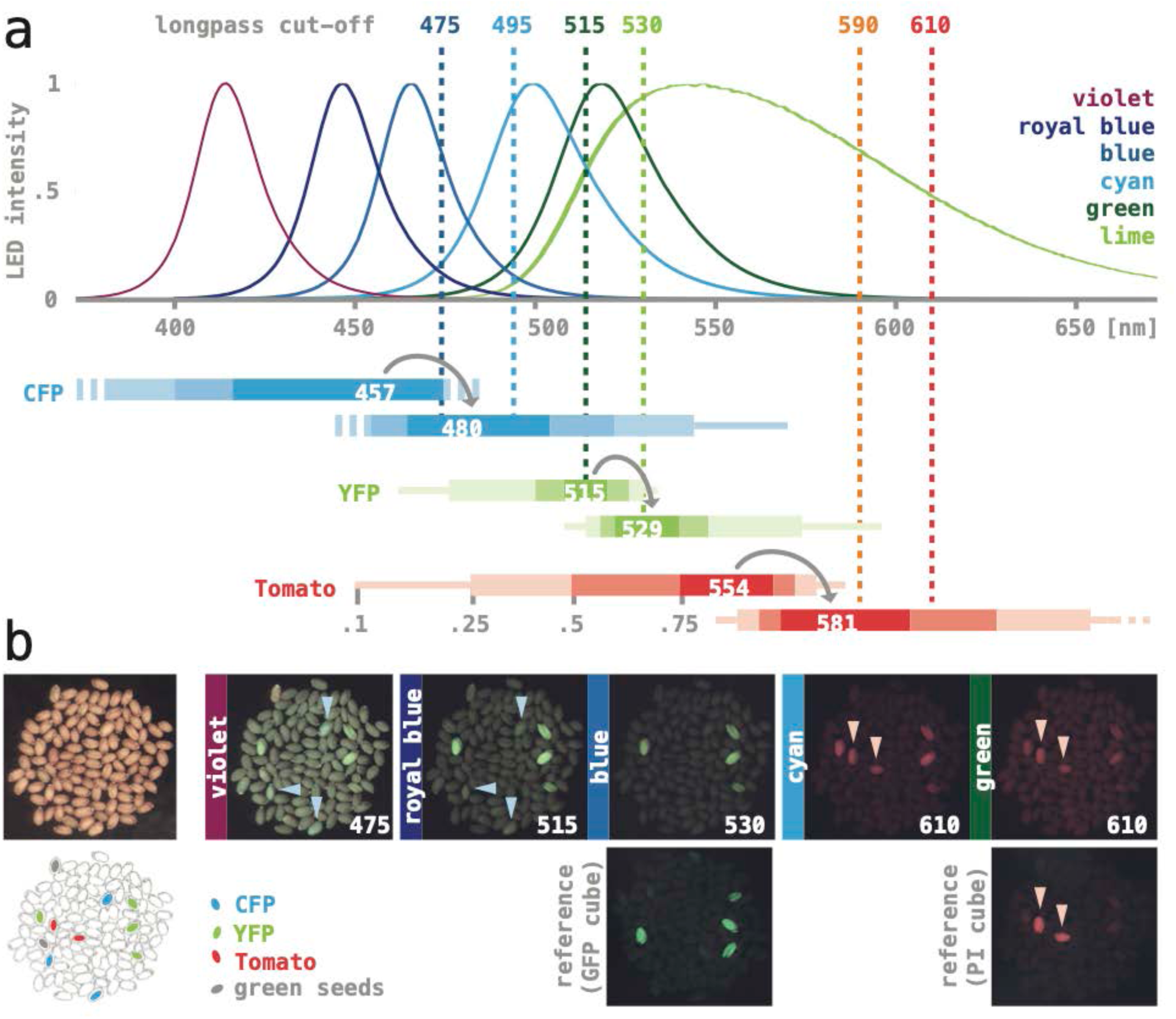
LED / coated glass filter illumination for imaging seed fluorescence. (**a**) Dotted lines mark the cut-off of longpass colored-glass alternative filters (Newport 20CGA-475, −495, −515, −530, −590, −610; values from www.newport.com). Curves show normalized emission spectra of five high-intensity LED assemblies (‘violet’: Luxeon Star SZ-05-U9, ‘royal-blue’: -H4; ‘blue’: -H3; ‘cyan’: -H2; ‘green’: -H1; ‘lime’: -H9; graphs adapted from www.luxeonstar.com). Horizontal bars represent the excitation (top) and emission spectra (bottom) of three fluorescent proteins below (maxima are listed; shaded intervals mark 0.75, 0.5, 0.25, and 0.1 of the respective maxima; values from www.fpbase.com; Lambert, 2019). (**b**) A sample of control seeds imaged with different illuminations. A bright field image and a trace of the seeds are shown on the left, with the position of CFP-, YFP-, and Tomato-expressing transgenics, as well as two greenish, chlorophyll-containing seeds highlighted. The LED assemblies and the cut-off of colored-glass alternative filters used to generate the images are noted on each panel. Images taken with the benchmark illumination are shown in the bottom row. Light-blue arrowheads in ‘475’ and ‘515’ point to CFP-expressing seeds, pink arrowheads in ‘610’ and the PI benchmark point to Tomato-expressing seeds. All fluorescence images were taken with the same camera settings except for exposure time, which was either 0.5 s (515, 530, GFP benchmark) or 1 s (others).

Seeds expressing YFP could be readily imaged using ‘royal blue’ or ‘blue’ LED lights (∼440 nm and ∼470 nm maximum) and 515 nm or 530 nm emission filters (Fig. 2b). These illuminations produced a brighter background compared to the benchmark, but the overall contrast remained high. CFP-expressing seeds were best detectable when illuminated with a ‘violet’ LED light (∼410 nm maximum) combined with a 475 nm emission filter, and Tomato-expressing seeds when illuminated with ‘cyan’ or ‘green’ LED lights (∼500 nm and ∼520 nm maximum) combined with 590 nm or 610 nm excitation filters. However, CFP- and Tomato-fluorescence was significantly dimmer, resulting in low signal-to-noise ratios (Fig. 2b); in addition, the illuminations were not selective: YFP-expressing seeds often appeared as bright as CFP- or Tomato-expressing seeds. Remnants of chlorophyll present in a small fraction of the seeds created relatively strong red fluorescence, in particular when viewed with ‘violet’ LED lights.

*Arabidopsis* is most commonly transformed by infiltrating live plants with *Agrobacterium* (Clough & Brent, 1998), and typically less than a percent of the seeds harvested from treated plants will be transgenic. As a stringent practical test, we used the benchmark instrument as well as a standard dissecting scope and LED / colored-glass alternative filter illumination to select primary transgenic, fluorescent seeds form the same sample collected after *Agrobacterium* infiltration.

For T-DNAs with a YFP marker, more than 80% of the YFP-fluorescent seeds detected with the benchmark instrument were also detected using illumination with LEDs and colored-glass alternative filters. For T-DNAs with a Tomato marker, the ratio was much lower (about 20%), implying that only transformation events resulting in strong Tomato-fluorescence can be reliably scored (our benchmark instrument was not fitted for imaging CFP, preventing a similar test with the CFP maker).

In summary, ‘royal-blue’ or ‘blue’ LED lights in combination with 515 nm or 530 nm longpass emission filters provide effective illumination for imaging YFP fluorescence in seeds. CFP and Tomato seed fluorescence can be detected using ‘violet’ and ‘cyan’ or ‘green’ LED lights combined with a 475 nm or 610 nm longpass emission filters, respectively – however, the sensitivity is comparatively low. Step-by-step instructions for assembling light sources as used here, including a supplier list and current prices of the components, can be found in the supporting material (S2 File).

### Targeting GRF genes individually: Cas9 expression with the polyubiquitin promoter results in ten-fold higher mutation rates than expression with the EC1 promoter

We chose the nine *Arabidopsis* GROWTH-REGULATING FACTORS (GRFs) as a test-case for CRISPR/Cas9 mutagenesis with our vectors. GRF genes are found in the genomes of streptophytes, a phylogenetic group including the land plants and their algal precursors, such as the charophytes. GRF proteins have two defining structural features: a QLQ domain with the invariant core of QX3LX2Q, and a WRC domain with a conserved sequence of three cysteine and one histidine residues (Fig.3 a; reviewed in Omidbakhshfard & al., 2015; Kim, 2019; it should be noted that the WRC domain does not conform to the consensus of C3H zinc-fingers, since the conserved positions show different spacing; Wang & al., 2008). GRFs are known to associate with GRF-INTERACTING FACTORs (GIFs) and to bind DNA in a sequence-specific manner. A phylogenetic analysis reveals that the *Arabidopsis* GRF genes fall into five clades, all of which were already present in the last common ancestor of flowering plants (Fig.3 b; Omidbakhshfard & al., 2015). GRF5/GRF6 and GRF7/GRF8 were separated in a whole genome triplication event at the base of the eudicots (∼130 million years ago; Jiao & al., 2012); GRF1/GRF2 and GRF3/GRF4 reside on large syntenic blocks (PPGD database, chibba.agtec.uga.edu/duplication; Lee & al., 2012) and were separated in the alpha whole genome duplication (∼30 million years ago, before the split of *Arabidopsis* and *Brassica*; Vision & al., 2000; Ermolaeva & al., 2003). Insertion alleles have been reported for all GRFs with exception of GRF6 (Fig 3 a); however, in many cases the insertion sites are in the promoter, in introns, or downstream of the WRC motif, and it is not clear that gene function is completely abolished. According to expression data in the public domain, GRF transcripts appear to accumulate predominantly in tissues with high mitotic rates, such as in the shoot apical meristem (SAM) and reproductive organs (Fig. 3 b; data taken from Klepikova & al., 2016; see Lee & al., 2018, for expression of GRF reporter genes in inflorescences).

**Fig 3.**
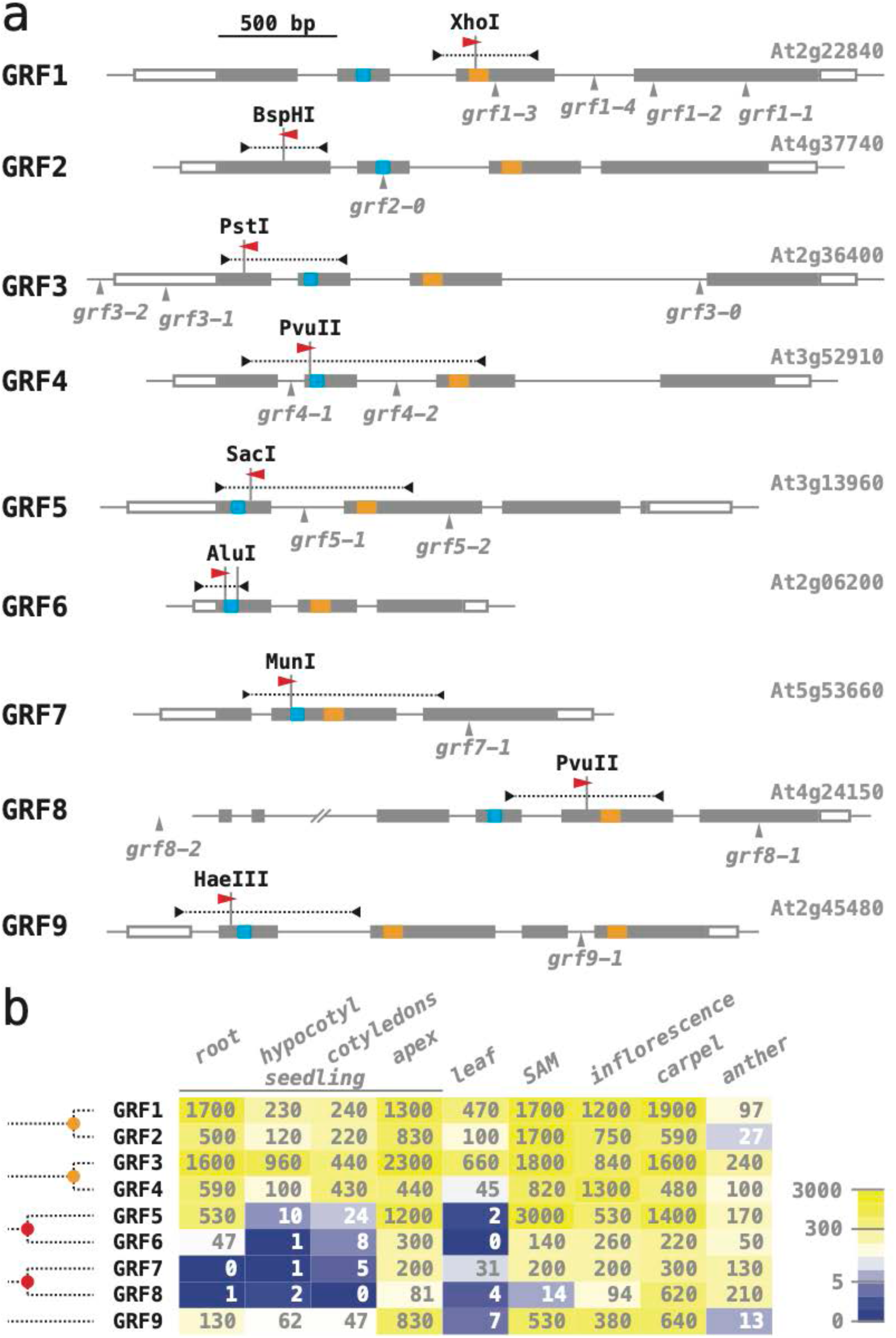
The GROWTH-REGULATING FACTOR (GRF) gene family of *Arabidopsis*. (**b**) Schematic organization of the *Arabidopsis* GRF genes (aligned at the translational start site). Exons are shown as boxes, with coding sequences filled in grey, the QLQ motif in blue, and the WRC motif in orange. The position and direction of protospacer sequences are indicated by a red arrowhead, with the targeted restriction sites sown above and the gene fragments that were amplified to screen for induced mutations as dotted lines. All previously reported GRF alleles are due to T-DNA insertions, mapped below the gene models; they are described in: Kim & al. (2003), *grf1-1, grf1-2, grf2-0, grf3-0*; Horiguchi & al. (2005), *grf5-1, grf9-1*; Kim & Lee (2006), *grf4-1*; Hewezi & al. (2012), *grf1-3, grf1-4, grf3-1, grf3-2*; Kim & al., (2012), *grf7-1, grf8-1, grf8-2*; Lee & al. (2018), *grf5-2*). (**b**) Overview of GRF mRNA expression (from travadb.org; Klepikova & al., 2016). Numbers represent the normalized average count per million reads; note that the scale of the color scheme is logarithmic. Samples of ‘seedlings’ were collected one day after germination; ‘apex’ represents the shoot meristem and surrounding tissues; ‘leaf’ represents the third rosette leaf at the time of flower opening; ‘SAM’ represents the vegetative shoot apical meristem 8 days after germination; ‘carpels’ and ‘anthers’ were harvested at the time of flower opening. Phylogenetic relationships of the *Arabidopsis* GRF genes are sketched on the right side (after Omidbakhshfard & al., 2015): the five GRF sub-clades found in *Arabidopsis* date back to before the last common ancestor of flowering plants; the alpha whole genome duplication event (∼30-35 million years ago) is marked by an orange dot, the gamma triplication event at the base of the eudicots (∼120 million years ago) by a red dot.

We followed two criteria for selecting specific spacer sequences targeting individual GRF genes from the CRISPR-Plant database (www.genome.arizona.edu/crispr2; Xie & al., 2014): first, the spacers had to target an exon upstream of the WRC motif, such that induced alleles would likely be nulls; second, the predicted Cas9 cut site had to lie within the recognition sequence of a restriction enzyme, such that induced mutations could be detected by PCR and restriction digest (Fig 3b; spacer sequences are listed in S3 File). Annealed oligonucleotides encoding the selected spacer sequences were then inserted into T-DNA vectors expressing Cas9 from either the EC1 or the UBI10 promoter. For each construct, ∼25 T1 seedlings were selected and their DNA was tested for mutant sectors; samples were scored as positive if ∼50% or more of the PCR product remained uncut after restriction digest (see Fig 4a for an example).

**Fig 4.**
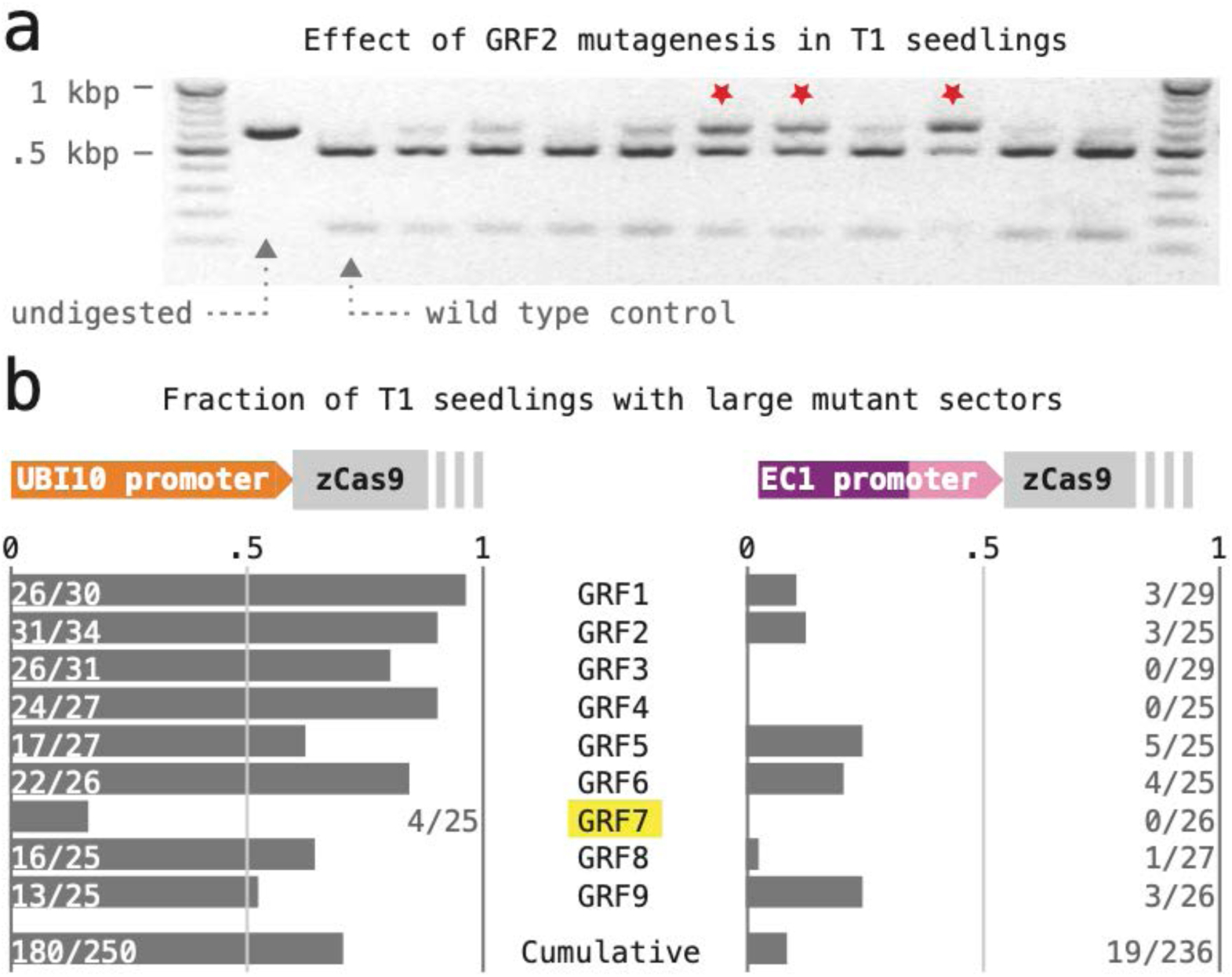
CRISPR/Cas9 mutagenesis with constructs expressing a single sgRNA. (**a**) The rate of induced mutations was estimated based on the occurrence of large mutant sectors in the T1 generation. Results for 11 T1 seedlings transformed with a construct expressing Cas9 from the EC1 promoter are shown as an example; the red stars mark cases where about half or more of the total DNA was undigested – these cases were scored as positive. (**b**) Cas9 apoprotein was expressed either from the polyubiquitin10 (UBQ10, left) or the egg cell-specific EC1 promoter (right). Estimated mutation rates are listed below.

The frequency of seedlings with large mutant sectors was taken as a proxy for the rate of CRISPR/Cas9-induced mutations at the nine GRF loci (Fig 4b). Our results reveal that expression of Cas9 with the polyubiquitin10 promoter caused almost ten-fold higher overall mutation rates than expression of Cas9 with the EC1 promoter (UBI10: 0.72, n=250; EC1: 0.058, n=236). By comparison, the mutation rates at different target loci (induced by different sgRNAs) showed much less variability: when Cas9 was expressed from the UBI10 promoter, frequencies of greater than 50% were obtained for all GRFs except GRF7. The GRF7 sgRNA also performed poorly when Cas9 was expressed from the EC1 promoter.

### A collection of reference null-alleles for the *Arabidopsis* GRF family

We next examined the germline transmission of CRISPR/Cas9-induced mutations (Fig 5). For each target locus, ∼10 T1 plants were grown to maturity and tested for mutant sectors using DNA extracted from rosette leaves or the primary inflorescence; sectored plants were allowed to self-fertilize and their seed harvested; ∼3-6 non-fluorescent, transgene-free T2 seed per positive T1 line were then propagated on soil, and the resulting plants assayed again. Despite this small sample size, mutant alleles were recovered in most GRF genes (GRF1: 3 alleles from 3 T1, testing 3 T2 each; GRF3 & GRF4: 2 alleles from 3 T1, testing 3 T2 each; GRF5 & GRF6: 3 alleles from 7 T1, testing 3 T2 each); only with GRF7 and GRF8 was it necessary to examine the progeny of more than 10 T1 plants.

**Fig 5.**
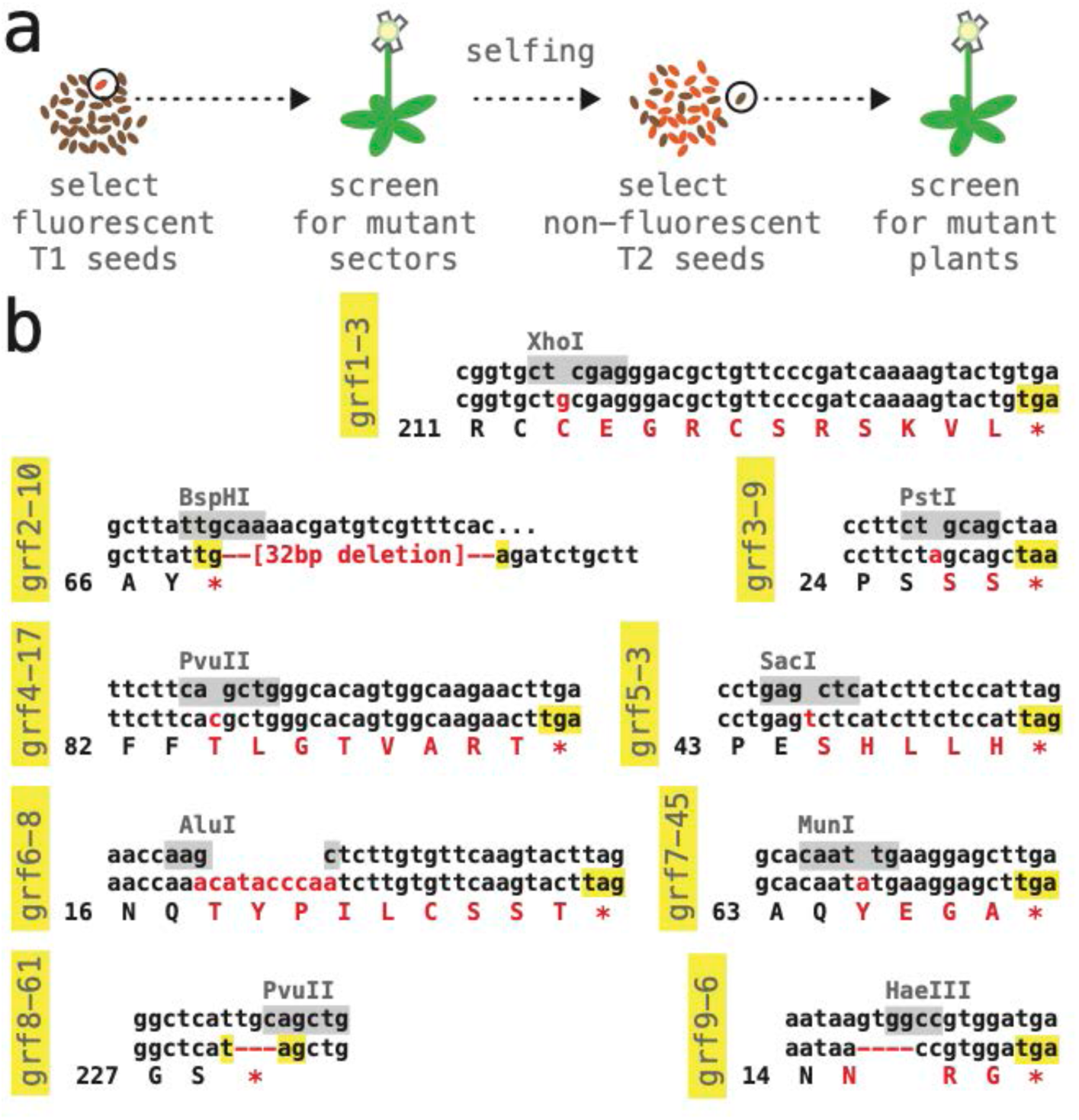
Collection of reference null alleles in *Arabidopsis* GRF genes. (**a**) Selection and counter-selection scheme for identifying CRISPR/Cas9-induced mutations; see text for details. (**b**) Molecular lesions of reference alleles. The wild type DNA sequence surrounding the CRISPR/Cas9 cut site is listed on top, with restriction site used to identify mutations highlighted in grey; the middle and bottom row show the DNA and predicted protein sequence of the mutant allele; inserted or deleted nucleotides as well as amino acid changes are shown in red. All alleles are predicted to cause a premature stop (highlighted in yellow).

Stably transmitted GRF alleles were also identified in the non-transgenic progeny of ∼10-20 T1 plants that had been bulk-harvested blindly, without screening for mutant sectors (GRF2: 3 alleles, testing 10 T2 from a pool of 10 T1; GRF8: 1 allele, testing 20 T2 from a pool of 10 T1; GRF9: 5 alleles, testing 20 T2 from a pool of 10 T1). While this simpler protocol may reduce time and effort in large-scale experiments, it is less well controlled and showed no substantial benefits in our case.

Sanger sequencing of the induced mutations revealed that the vast majority were due to small insertion/deletion events at the predicted CRISPR/Cas9 cut sites. From this collection, we selected *bona-fide* null alleles in all nine *Arabidopsis* GRF genes (Fig 5b). Homozygous single mutant plants could be obtained with all these reference alleles and showed no obvious abnormalities. Seed samples are available from the *Arabidopsis* stock center (see Methods).

### Targeting pairs of GRF genes: similar mutation rates were obtained by expressing two sgRNAs as independent genes or as part of one polycistronic gene including a tRNA buffer

Plants simultaneously expressing more than one sgRNA species have been produced by placing independent sgRNA genes on a single T-DNA or, alternatively, by constructing a polycistronic transcription unit, in which segments encoding sgRNAs alternate with segments encoding a tRNA; the cellular tRNA processing machinery will excise these tRNA segments post-transcriptionally, liberating the sgRNAs (Xie & al., 2015). We have directly compared the efficiency of these two designs by generating T-DNAs targeting pairs of GRF genes (GRF1/2, GRF3/4, GRF5/6, and GRF7/8) either with sgRNAs expressed form separate promoters (U2-26, At3g13855; U6-29, At5g46315) or with sgRNAs derived from a polycistronic transcript (Fig 6). DNA fragments encoding the respective combinations were produced by PCR with primer combinations that included the gene-specific spacer sequences as well as terminal *BsaI* sites (as in Xing, & al., 2014; a plasmid containing the U6-29 promoter and a synthetic DNA fragment containing an alanine tRNA sequence served as templates; see Methods and S3 File for details). The sgRNAs contained the same gene-specific spacer sequences as previously, and all constructs were based on vectors driving Cas9 expression from the UBI10 promoter.

**Fig 6.**
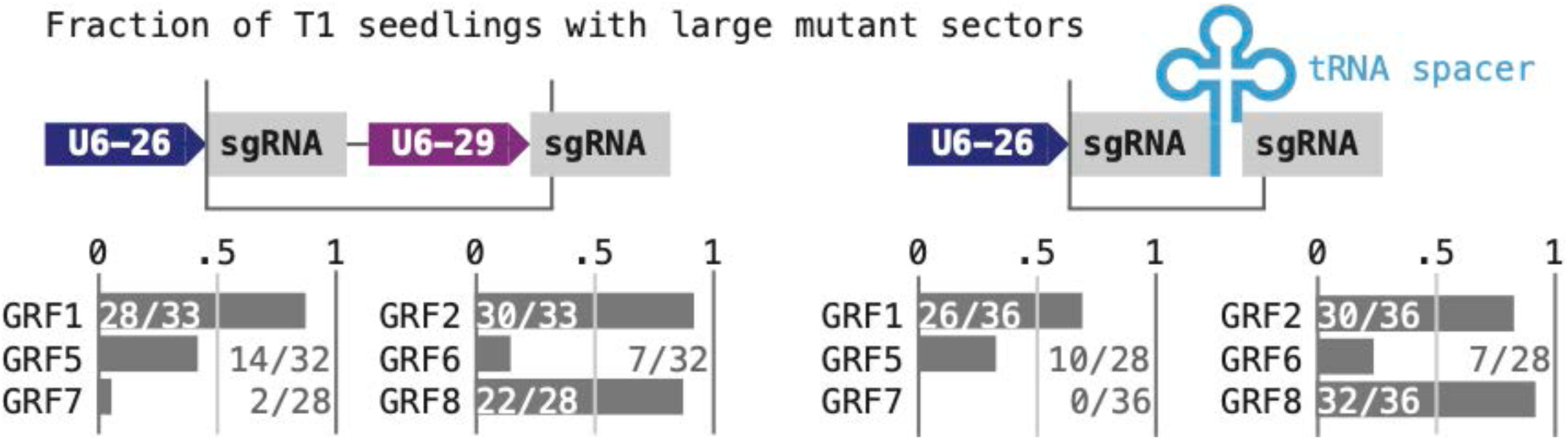
CRISPR/Cas9 mutagenesis with constructs expressing two sgRNAs. Different combinations of two sgRNA species were expressed either from separate genes (top left) or from a polycistronic gene with a tRNA buffer (top right); DNA fragments inserted into the *BsaI* cloning site of the vector boxed; ‘U6-26’ and ‘U6-29’, small RNA promoters. Estimates of the mutation rates at target genes are shown below, with the targets listed as the sgRNAs were arranged on the constructs.

Two attempts at transforming T-DNAs targeting GRF3/4 failed to produce fluorescent T1 seeds, suggesting that this combination of sgRNAs may be detrimental to transformed cells. T1 seedlings harboring the remaining three pairs of constructs were assayed for large mutant sectors as before. Averaged over all target genes, the mutation rates in this experiment seemed slightly lower than the rates observed with single sgRNAs; but they were not dependent on how the sgRNA species were generated (Fig 6; independent promoters: 0.55, n=186; separated by tRNA: 0.52, n=200; compared to 0.72, n=250, when targeted individually), nor was there an apparent correlation of the mutation rate to the position of an sgRNA on the construct (GRF1, GRF5, GRF7: 0.57, n=82, when targeted individually; 0.41, n=193, when targeted as part of a pair; GRF2, GRF6, GRF8: 0.81, n=85, when targeted individually; 0.61, n=118, when targeted as part of a pair). Loci showing the highest mutation rates when targeted individually also showed the highest rates when targeted as part of a pair. Interestingly, mutations in the two members of a pair did not arise independently: the large majority of all T1 seedlings either tested wild type at both target genes (55) or harbored mutant sectors in both (70); the remaining seedlings showed large sectors only at the target with a higher overall mutation rate (68; n=193, combined data for both types of constructs). These findings imply that CRISPR/Cas9 activity in our experiment varied more substantially between transgenic lines than it did between complexes containing different sgRNA species.

### Simultaneous targeting of the entire gene GRF family: multiplexing of four or more CRISPR RNAs results in vastly different mutation rates at different target genes

Finally, we explored multiplexing sgRNAs as a means of creating multiple mutant combinations in a moderate number of target loci. To minimize repetitive sequences in the constructs, we combined the multiplexing approaches assessed above: two small RNA genes driven by a U6-26 and a U6-29 promoter, respectively, were arranged in tandem; each gene produced a poly-cistronic transcript encoding two sgRNAs separated by an alanine tRNA (Fig 7a). Construct ‘1256’ had a Tomato selectable marker and targeted GRF1, GRF2, GRF5, and GRF6; construct ‘3478’ had a YFP selectable marker and targeted GRF3, GRF4, GRF7 and GRF8 (see Methods and S3 File for details).

**Fig 7.**
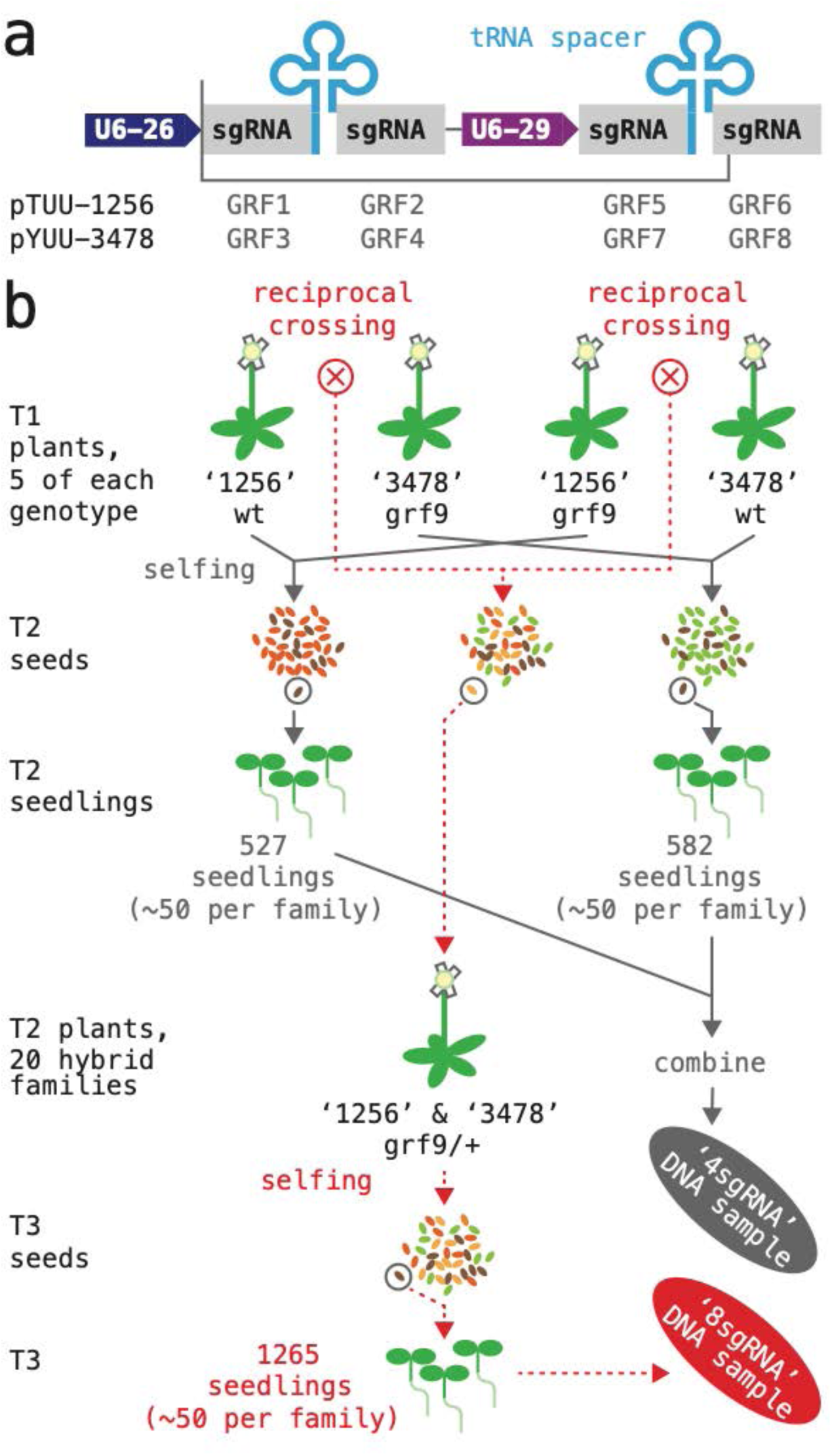
Simultaneous CRISPR/Cas9 mutagenesis of four and eight GRF genes. (**a**) A schematic of the two T-DNA constructs expressing four sgRNA species are shown on top; DNA fragments inserted into the *BsaI* cloning site of the vector boxed; ‘U6-26 and U6-29’, small RNA promoters. (**b**) Flow-chart illustrating how the two DNA samples for amplicon sequencing were generated. The ‘4sgRNA’ sample (grey arrows) represents plants that had been subjected to mutagenesis by either the ‘1256’ or the ‘3478’ construct; for each construct, 10 T1 plants were grown to maturity and harvested; ∼50 transgene-free seeds per T1 plant were germinated and combined for DNA extraction. The ‘8sgRNA’ sample (dotted red lines) represents plants that had been subjected to mutagenesis by both constructs; reciprocal crosses between pairs of ‘1256’ and ‘3478’ T1 plants were performed (total of 20 crosses), and T2/F1 plants containing both constructs selected; ∼50 transgene-free seeds per T3/F2 family were germinated and combined for DNA extraction.

The two T-DNAs were used to generate populations of plants in which either four or eight GRF genes were being mutagenized; in addition, the plants were segregating for the *grf9-6* reference allele (GRF9 was not targeted by the any of the T-DNAs; Fig 7b). Toward this, both constructs were transformed separately into wild type as well as *grf-9-6* mutant plants. For each combination, five T1 plants that contained a large mutant sector in at least one of the target genes were then selected, allowed to self-pollinate, and harvested – yielding 20 families of T2 seeds. In addition, reciprocal crosses were performed between the five wild type plants harboring the ‘1256’ transgene and the five *grf9-6* plants harboring the ‘3478’ transgene (total of 10 crosses), as well as the five *grf9-6* plants harboring the ‘1256’ transgene and the five wild type plants harboring the in the ‘3478’ transgene (total of 10 crosses). From each cross, ∼5 T2 / F1 seed showing Tomato-as well as YFP-fluorescence were propagated; the resulting plants (*grf9-6*/+, hemizygous for both constructs, representing 20 independent transformation events) were allowed to self-pollinate and harvested – yielding 20 families of T3 seeds.

The frequency of GRF mutations induced by the ‘1256’ or ‘3478’ constructs either separately or in combination was then estimated by amplicon-sequencing. Two samples of pooled DNA were prepared for this purpose: the ‘4sgRNA’ sample represented a population in which four genes were targeted simultaneously (grey in Fig 7b; 527 and 582 seedlings produced by T1 plants harboring ‘1256’ and ‘3478’, respectively; ∼50 seedlings per family; half of the T1 parents were *grf9-6*); the ‘8sgRNA’ sample represented a population in which eight genes were targeted simultaneously (red in Fig 7a; 1265 seedlings total, ∼50 per family; all T2 parents were *grf9-6*/+). Selection of non-fluorescent seeds ensured that, barring sampling errors, our samples only contained germline-transmitted mutations. A small number of seedlings with known mutations in GRF9 was added to both samples for control purposes (see Methods).

DNA fragments covering the predicted CRISPR/Cas9 cut sites at all nine GRF genes were generated by PCR, barcoded, and sequenced on an Illumina platform (see Methods and S3 File for details). On average, >95% of the resulting reads mapped to their amplicon, and the number of reads expected to be generated by one GRF allele of one seedling in the sample was ∼46 (between ∼23 and ∼56; see Methods for summary statistic and representation of controls included in the samples). We used the AGEseq tool (Analysis of Genome Editing by sequencing; Xue & Tsai, 2015) to determine the frequency of small deletions or insertions in the DNA samples (Fig 8; transitions and transversions were not considered, since they are not commonly induced by CRISPR/Cas9 and can be difficult to distinguish form PCR or sequencing artifacts; deletions of ∼100 bp or greater, which are more common, would have escaped our analysis). Nearly all lesions mapped exactly to the predicted CRISPR/Cas9 cut site; in the remaining cases (grey dots in Fig 8), the mutations mapped only one to two positions off. The most common types of lesion were single base pair insertions, and deletions of one or two base pairs. Five relatively long insertions, ranging from 15 to 55 bp, were also represented in our collection (Fig 8); while two of these insertions originated from the mutagenized GRF locus, we were unable to determine the origins of the remaining three). Although different target genes showed slightly different spectra of lesions, we saw no evidence for microhomology-based repair at the CRISPR/Cas9 cut site (as discussed in Vu & al., 2017; Sfeir & Symington, 2015).

**Fig 8:**
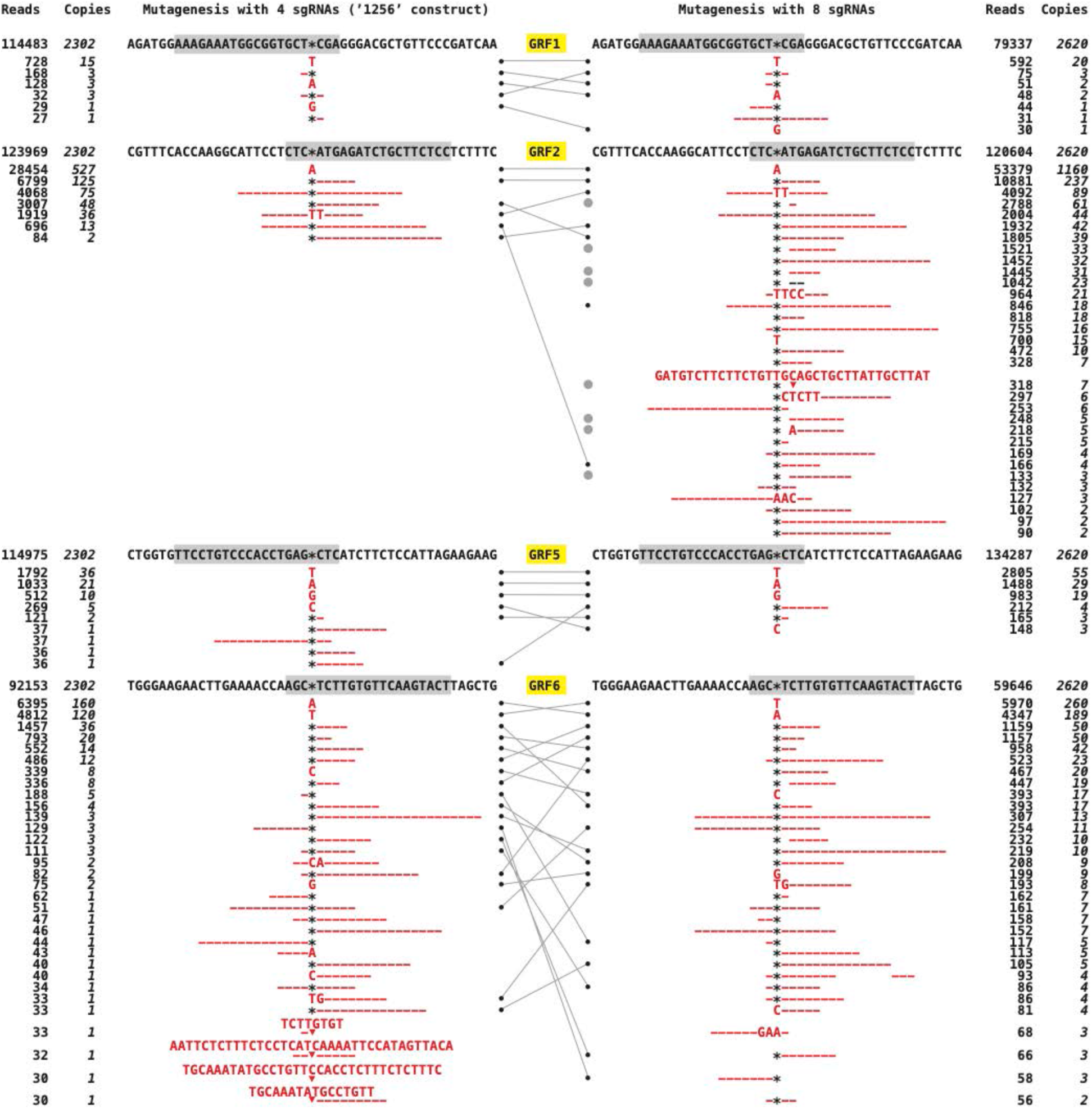
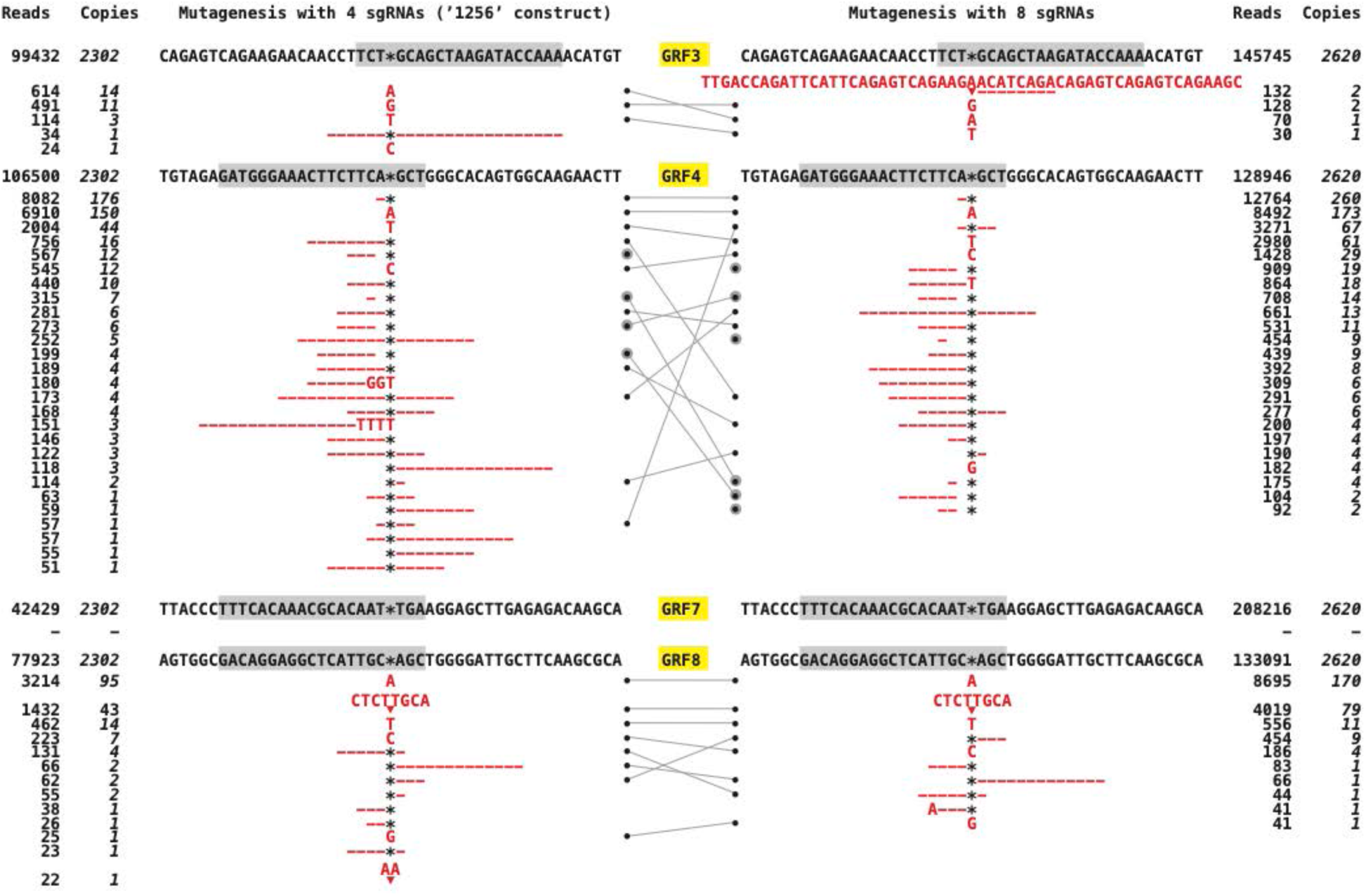
Spectrum of CRISRP/Cas9-induced mutations at different target genes. Spectrum and frequency of insertion-deletion events identified by AGEseq in GRF target genes. Targets are arranged according to their position on the ‘1256’ or ‘3478’ constructs, protospacer sequences are highlighted in grey, and the predicted Cas9 cut site is represented by a star. The total number of mapped reads supporting an allele is listed under ‘reads’, and the estimated number of seedlings in the pool that are heterozygous for the allele is shown in cursive under ‘copies’ (see Methods). Alleles that were found in both DNA pools are marked with a black dot and connected by grey lines; these alleles may not have been induced independently, since the same T1 plants gave rise to both pools. Grey dots mark alleles in which the predicted CRISPR/Cas9 cut site is not part of the lesion. The 35 bp insertion in GRF2 (GATGTC…) and the 16 bp insertions in GRF6 two alleles (TGAAAA…) originate from within the gene.

The overall effects of the two constructs did not vary much between our two populations: the target genes of ‘1256’ were mutated with an average frequency of ∼32% when the construct was acting by itself and ∼29% when acting in combination with ‘3478’; for the ‘3478’construct, the corresponding values were ∼7% and ∼10%. This observation implies an additive interaction on average (possibly because the average ratio of Cas9 apoprotein to total sgRNA remains constant). The perhaps most striking result of the experiment was that mutation frequencies at individual target genes were vastly different and often strongly affected by multiplexing. A compilation of the estimated mutation rates gathered in all our experiments reveals that multiplexing sgRNAs drastically reduces the frequency of CRISPR/Cas9-induced lesions at some target genes, while showing only small effects at others (Fig 9; rates were normalized to GRF2, which showed the highest values in all experiments; see legend for details). When tested individually, all sgRNAs (except for the one targeting GRF7) were roughly similarly effective, producing large mutant sectors in greater than 50% of the T1 seedlings. The GRF2 sgRNA continued to induce mutations at consistently high rates: 95% by itself; 87% in combination with a GRF1 sgRNA; 73% and 75% when combined with three or seven other sgRNA species, respectively. In contrast, the mutation rates associated with the majority of sgRNAs show a more or less steep downward trajectory with increasing number of total sgRNA species, implying that multiplexing successively and more or less drastically reduces the activity of these sgRNAs.

**Fig 9:**
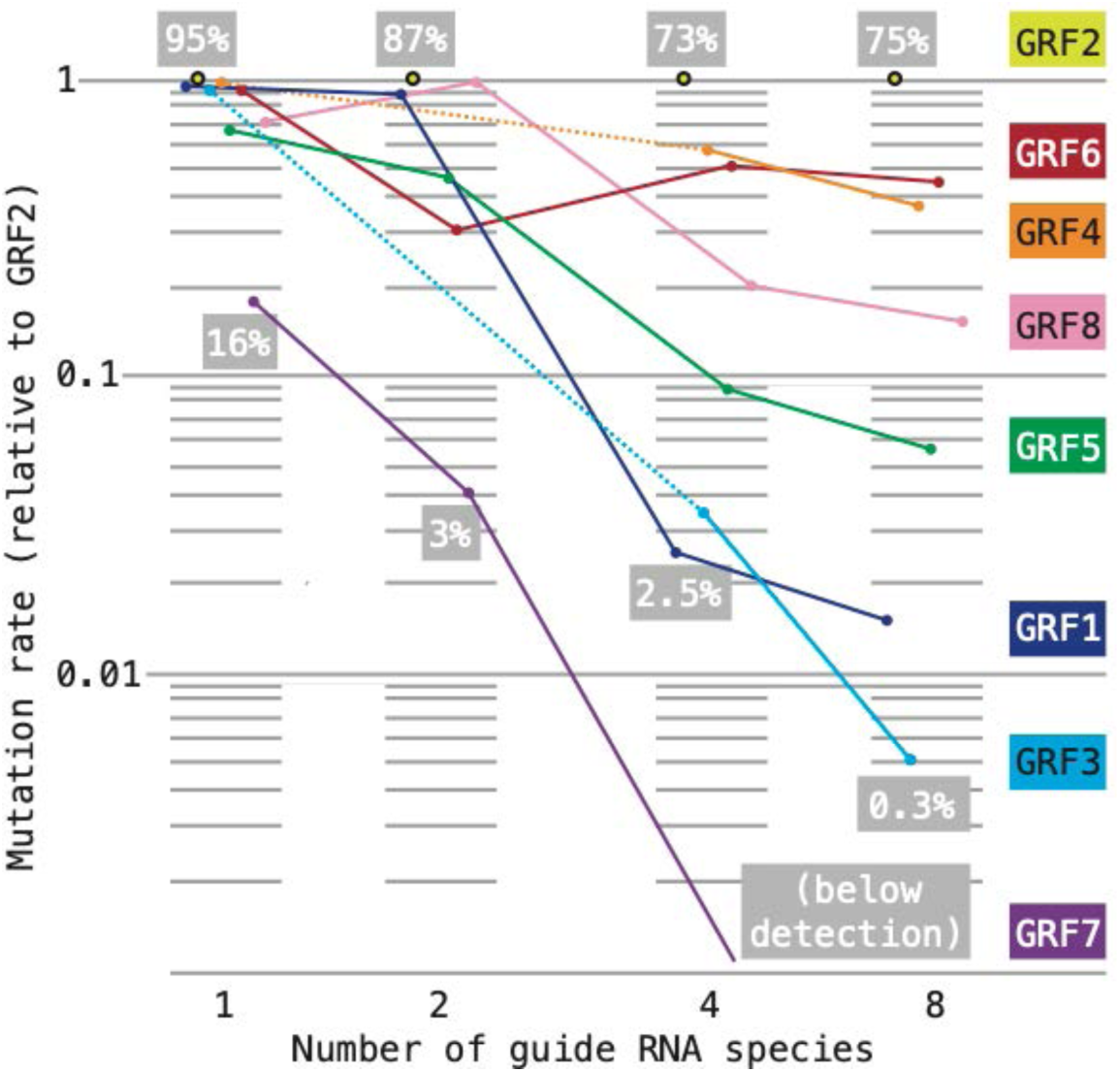
Effect of sgRNA multiplexing on mutation rates at different target loci. Our estimates of mutation rates are not directly comparable: in experiments with one or two sgRNAs, we assessed the frequency of large mutant sectors in ∼24 T1 seedlings; in experiments with four or eight sgRNAs, we assessed the frequency of germline-transmitted mutant alleles in ∼1000 seedlings. Throughout, the most effective sgRNA was the one targeting GRF2. For the purpose of comparison, the mutation rates estimated in each experiment were normalized to GRF2 (set to ‘1’ on the y-axis; note that the scale is logarithmic); the number of sgRNA species in the experiment is listed on the x-axis. To convey the range of mutation frequencies within each experiment, the highest and lowest observed values are listed in the grey boxes (expressed as percentage of the total sample).

## Discussion

A large variety of binary vectors is available for targeted mutagenesis in plants. Here, we add vectors enabling selection and counter-selection of transgenes on the basis of seed bio-fluorescence to the toolbox of CRISPR/Cas9 plasmids created by Xing & al. (2014) and Wang & al. (2015). The value of fluorescent markers for removing CRISPR/Cas9 transgenes once they are no longer needed has been recognized previously (Gao & al., 2016; Wu & al., 2018); however, the T-DNAs described by these groups were designed for large-scale experiments and rely on Gibson assembly for a high-throughput construction pipeline. In contrast, the vectors we describe here are suited for small-scale projects and minimal lab settings: construction relies on standard molecular cloning routines that require no specialized equipment or reagents; *BsaI*-cut vector fragments may be prepared in advance and stocked; plasmids expressing one sgRNA are generated by combining the vector with annealed synthetic oligonucleotides – as rapid, simple, and cheap a cloning procedure as we know; finally, transgenic seeds can be easily identified by illumination with inexpensive LED lights and colored-glass alternative filters. We tested these tools on the *Arabidopsis* GRF genes and recovered new reference alleles with frame-shift mutations truncating the predicted protein sequences before or within the conserved WRC domain for all nine members of the family with small effort (∼30 PCR-based tests in most cases). With exception of the GRF7 sgRNA, all guides caused high rates of induced mutations when expressed individually (although they were selected from a large database without regard to sequence composition or secondary structure).

A convenient method for simultaneously mutagenizing multiple loci would greatly simplify the genetic analysis of functional redundancies, and several platforms for multiplexing CRISPR RNAs in plants have been developed with this aim in mind (reviewed in Najera & al., 2019). Ma and colleagues (2015) simultaneously targeted eight rice FT-like genes and recovered primary transgenic seedlings with mutations in seven of the targets. These multiple mutant plants showed visible phenotypes consistent with a loss of FT-like function, but it was not reported whether the induced mutations were transmitted to the next generation. Xie and colleagues (2015) simultaneously targeted five rice MAP kinase genes and found that ∼50% of the primary transgenics harbored editing events in all targets (again, it was not reported whether the mutations were heritable). Following similar overall procedures as we, Zhang and colleagues (2016) targeted six *Arabidopsis* PYR1-like genes and identified one primary transgenic plant (out of 15) that contained mutations in all of them; importantly, the mutations were germline-transmitted and enabled construction of a sextuple mutant.

In contrast to these studies, our experiments enabled us to trace the trajectory of mutation rates at different GRF targets in plants expressing a single, two, four and eight sgRNA species. The results revealed strong interference effects: while some targets always showed lesions with a high frequency, the mutation rate at most targets dropped, often drastically, as the number of sgRNA species increased. This effect could be caused by synthetic lethality or similarly strong interactions between induced alleles. Such interactions are often seen with mutations in closely related genes that can provide overlapping function. Phylogeny indicates that the gene pairs GRF1/GRF2, GRF3/GRF4, GRF5/GRF6, and GRF7/GRF8 arose in duplication events within flowering plant lineage (Fig. 3), and it does not seem unlikely that the two genes of a pair retain similar activities. Multiplexing appeared to affect the two members of a pair in markedly different ways: while one of the genes (GRF2, GRF4, GRF6, GRF8) retained relatively high mutation rates, rates dropped sharply in the other (GRF1, GRF3, GRF5, GRF7; Fig. 9). This apparent dichotomy is consistent with the idea that double mutant gametophytes or sporophytes are impaired or inviable. Our constructs simultaneously targeting two GRF genes revealed that CRISPR/Cas9-induced mutations do not arise independently; rather, it appears that mutations induced by the less effective sgRNA predominantly arise concomitantly with mutations induced by the more effective sgRNA. Since we assessed apparent mutation rates at the seedling stage, this asymmetry would explain how the loss of double mutant genotypes due to synthetic lethality may affect one gene of a pair much more dramatically than in the other. On the other hand, a survey of more than 1000 individuals derived from plants that were harboring the ‘1256’ as well as ‘3478’ constructs and were heterozygous for the *grf9-6* reference allele (the same plants that gave rise to the seedlings analyzed in your amplicon sequencing experiment) uncovered only a single line segregating for embryo-lethality and no lines with an obviously defective female gametophyte (male gametophytes were not examined). Similarly, multiple mutants of previously described GRF insertion alleles show only mild defects (Kim & al., 2003; Horiguchi et al., 2005; Kim & Lee, 2006; Hezewi & al., 2012; Lee & al., 2018), such that there is little evidence for pervasive and strong detrimental interactions between GRF mutations.

A second factor influencing the trajectory of mutation rates appears to be inherent to the CRISPR/Cas9 machinery: multiplexing tends to amplify relatively small differences in the baseline activity of individual sgRNAs (assessed in the absence of other sgRNA species). An analogous, although less pronounced interference-effect upon multiplexing was observed in rice protoplasts and attributed to competition of sgRNAs for a limiting amount of Cas9 apoprotein (Xie & al., 2015). Any such competition would be exacerbated in our case, as multiplexing likely increases the relative abundance of sgRNAs with respect to Cas9 apoprotein: constructs targeting two or four GRF genes can potentially produce two or four times the amount of sgRNAs compared to constructs targeting a single gene (assuming similar activity of the two U6 promoters and no substantial losses in transcript processing). The idea that Cas9 abundance often limits genome editing events in plants is not new. Osakabe and colleagues (2016) reported that the accumulation of Cas9 apoprotein varied widely between independent transgenic lines and was well-correlated with the occurrence of induced mutations. Indeed, this effect is so strong that it has prompted the incorporation of ‘target-proxies’, fluorescent markers that include a target mimic, into CRISPR/Cas T-DNAs as a means of selecting transformation events with robust CRISPR/Cas9 activity (Li & al., 2020). Finally, it is well documented that the promoter driving Cas9 expression strongly affects the frequency of editing events (reviewed in Ma & al., 2016; Soyars & al., 2018; we too found tenfold lower mutation rates with the EC1 promoter than the UBI10 promoter).

Competition for a limiting amount of Cas9 would further imply that different sgRNAs associate with Cas9 apoprotein more or less effectively. The assembly of active CRISPR/Cas9 complexes is accompanied by significant changes in the conformation of Cas9 apoprotein (reviewed in Jiang & Dounda, 2017), but the role of the spacer sequence in this process (the only segment that differs between different sgRNA species) has so far not been described. We did not notice any obvious hallmarks in the primary sequence, GC-content, melting temperature, or predicted secondary structure of the GRF guide sequences that seem to correlate with their activity in our multiplexing experiments. Thus, competition effects as suggested by our results may be difficult to manage in practice and could pose potentially severe obstacles to multiplexing strategies.

## Materials & Methods

### Seed stocks, plant growth & transformation

The Columbia accession of *Arabidopsis thaliana* served as a wild type strain for transformation and CRISPR/Cas9 mutagenesis. Plants were grown under constant fluorescent light at ∼25°C on commercial potting mix (RediEarth, Sun Gro Horticulture) with slow-release fertilizer (Osmocote). For germination in sterile culture, seeds were surface sterilized in 70% ethanol for 1 minute, rinsed twice in 96% ethanol, briefly air dried, and transferred to plates containing 1% sucrose, 1% agar (Sigma A-1296) and 0.5x MS salts (MP Biomedicals 2623022). The GV3101 strain of *Agrobacterium* was used for plant transformation following the floral dip protocol (Clough & Brent, 1998). Reference alleles created as part of this study can be obtained from the *Arabidopsis* stock center (abrc.osu.edu): *grf1-3*, CS72426; *grf2-10*, CS72427; *grf3-9*, CS72428; *grf4-17*, CS72429; *grf5-3*, CS72430; *grf6-9*, CS72431; *grf7-45*, CS72432; *grf8-61*, CS72433; *grf9-6*, CS72434.

### Imaging

Seed fluorescence was imaged with an Olympus SZX12 stereo-microscope equipped with a Moticam 3.0 plus digital camera and a Kramer Scientific Quad internal illumination module connected to an X-cite 120 mercury lamp; filter sets for GFP (Kramer Scientific 184, with narrow band emission) and propidium iodide (Kramer Scientific 816) were used for YFP- and Tomato-fluorescence, respectively. The instrument did not have appropriate filters for imaging CFP-fluorescence. The improvised LED lamps we assessed as a low-cost alternative illumination are documented in S1 File.

### Plasmid construction

The CRISPR/Cas9 T-DNAs vectors enabling selection and counter-selection on the basis of seed fluorescence used in this study are available from Addgene (addgene.org): pCEE, TBA; pYEE, TBA; pTEE, TBA; pCUU, TBA; pYUU, TBA; pTUU, TBA.

T-DNAs expressing a single or two sgRNA were generated by conventional cloning (as described by Xing & al. 2014). T-DNA vectors were linearized by restriction with *BsaI* (we found that incubation overnight at 45°C gave best results) and gel purified. Vectors to be used in ligations with PCR products were treated with shrimp alkaline phosphatase (ThermoFisher 78390) prior to gel purification to remove their terminal phosphates. For single-sgRNA constructs, two non-phosphorylated complementary oligonucleotides encoding the 20 nt gene-specific spacer sequences plus appropriate overlapping ends for cloning (S2 File) were mixed (50 µM each, 2X SSC buffer) and annealed in a temperature gradient (96°C – 20°C); 1 µL of the annealing reaction was combined with ∼100 ng vector fragment (with intact 5’-phosphate ends) and ligated with T4 ligase (1 h at RT).

For two-sgRNA constructs, fragments containing the gene-specific spacer sequences, the sgRNA backbone and either a U6-29 promoter or a tRNA buffer were produced by four-primer PCR (described in Xing & al., 2014) with a proofreading enzyme (Q5 polymerase, New England Biolabs, M0491). Briefly, the reactions contained a pair of inner and outer primers at a ratio of 1:20 (50 nM and 1 µM, respectively); inner primers were designed to anneal to template plasmids and also encoded the spacer portion of the sgRNAs; outer primers were designed to use the DNA fragments produced by the inner primers during the first few PCR cycles as template and also contained sequences required for generating vector-compatible ends by *BsaI* digestion; in this way, the length of all primers could be limited to ∼40nt (S2 File). Two plasmids were used as templates: pGEM-2t, derived form a custom synthetic fragment (Integrated Gene Technologies, IDT) and containing a sgRNA backbone as well as an alanine tRNA spacer (see S3 File for an annotated sequence listing; the plasmid has been deposited with Addgene, TBA); and pCBC-DT1DT2 (Xing & al., 2014; Addgene #50590), containing a sgRNA backbone as well as a U6-29 promoter. ∼50 ng of *BsaI*-digested, gel-purified PCR products were ligated with ∼100 ng de-phosphorylated vector fragment (16°C, overnight).

T-DNAs expressing four sgRNAs were assembled from BsaI-linearized vectors and three PCR fragments using an NEBuilder kit (E2621, New England Biolabs). Appropriate PCR fragments were generated in three- or four-primer reactions similar to the ones described above, with the outside primer containing the 23-25 nt overlaps required for the assembly reaction (S2 File). The sgRNA genes of all constructs were verified by Sanger sequencing.

### Detection & sequencing of mutant alleles

Spacer sequences were selected such that CRISPR/Cas9 would cause a double strand break within the recognition sequence of a restriction enzyme, enabling the detection of induced mutations by PCR-based markers (S2 File). The same primers were used to amplify germline-transmitted mutant alleles for Sanger sequencing (with internal or PCR primers; S2 File).

### Amplicon sequencing & data analysis

Two samples of DNA of were generated for the purpose of mass-sequencing GRF mutations: the ‘4-sgRNA’ sample was extracted from 1151 T2 seedlings: 527 mutagenized with the ‘1256’ construct (230 from 5 wild type parents, 297 from 5 *grf9-6* parents), 582 T2 mutagenized with the ‘3478’ construct (297 from 5 wild type, 285 from 5 *grf-6* parents), and 42 controls (described below); the ‘8-sgRNA’ sample was extracted from 1310 T3 seedlings: 1265 mutagenized with both constructs (form 20 families of *grf9-6*/+ parents), and 51 controls. 10 independent transformation events of each construct are represented in this population (see Fig. 7 for pedigree). From each T1 parent or T2 family, ∼50 non fluorescent seeds were selected and grown on plate for 7 days; germinated seedlings were tallied and then combined to generate the two samples; the sample material was ground in liquid nitrogen and the DNA extracted following a modified CTAB protocol (Murray & Thompson, 1980).

For each GRF gene, ∼200 bp amplicons that included the CRISPR/Cas9 target sites were generated using non-proofreading Taq polymerase; the PCR primers contained tails for library construction (see S2 File for details). Amplicons were barcoded and sequenced on an Illumina platform at the UGA Georgia Genomics and Bioinformatics Core (dna.uga.edu). The resulting reads were aligned to a 60 nt wild type reference sequence centered around the predicted CRISPR/Cas9 cut site and analyzed for insertion and/or deletion events using AGEseq (Xue & Tsai, 2015).

GRF9 was not mutagenized in the experiment, such that the amplicon could be used to assess representation. Both DNA samples included a known number of GRF9 mutant seedlings for this purpose. 51% of the seedlings represented in ‘4sgRNA’ were from *grf9-6* parents (582 / 1151), and 51% of the mapped GRF9 amplicon reads contained the *grf9-6* mutation; similarly, 97% of the seedlings represented in ‘8sgRNA’ were *grf6-1/+* (1265 / 1310), and 50% of all mapped reads contained the *grf-6* mutation. In addition, the samples contained 24, 12, and 6 seedlings from *grf9-3/grf9-4, grf9-1/grf9-2*, and *grf9-7/grf9-8* parents, respectively. The *grf9-2* allele is a ∼180 bp deletion-insertion event that could not be amplified with our primers; *grf9-3, grf9-4, grf9-7*, and *grf9-8* harbor small deletions flanking the CRISPR/Cas9 cut site (indicated by a star): ttg*------atg, gtg-* atg, gtg-*ccg, tgg*-cgt, respectively; *grf9-1* contains a 9 bp insertion (upper case letters): tgg*AGTTTCGGAgga. The representation of these alleles in the two DNA samples is summarized in Table 1.

**Table 1.** Representation of *grf9* alleles in the sequences.

|  | Seedlings | Reads | Reads per copy<sup>1</sup> |
| --- | --- | --- | --- |
| **'4sgRNA' sample** |  |  |  |
| All | 1151 | 66145 | Expected: 29 |
| *grf9-3* & *grf9-4* | 24 | 1411 | 29 |
| *grf9-1* | 12 | 567 | 47 |
| *grf9-7* & *grf9-8* | 6 | 413 | 34 |
|  |  | Average: | 37 ± 9 |
| **'8sgRNA' sample** |  |  |  |
| All | 1310 | 60496 | Expected: 23 |
| *grf9-3* & *grf9-4* | 24 | 1268 | 26 |
| *grf9-1* | 12 | 567 | 48 |
| *grf9-7* & *grf9-8* | 6 | 413 | 39 |
|  |  | Average: | 38 ± 11 |

<sup>1</sup> The number of reads generated by one allele of one seedling in the sample; the expected value is all mapped reads divided by two-times the number of seedlings in the sample. The
observed values are the number of mutant reads divided by two-times the number of seedlings harboring the mutant alleles; an exception is *grf9-1*, which was calculated as number of mutant reads divided by the number of seedlings from *grf9-1/grf9-2* parents (since *grf9-2* cannot be amplified by our primers). Average ± standard deviation of the three observed values are indicated for both samples.

AGEseq reported 243 distinct insertion-deletion events in the sequences of GRF1-GRF8 amplicons (see Table 2 for a summary statistic and Fig 8 for details). We discarded 30 of these events as likely artifacts, since they were supported by fewer than 50% of the reads expected to be generated by one allele of one seedling (the representation of the *grf9* controls showed a standard deviation of ∼25%, suggesting that this cut-off is inclusive).

**Table 2.** Summary statistic of GRF1–GRF8 amplicon sequencing.

|  | GRF1 | GRF2 | GRF3 | GRF4 | GRF5 | GRF6 | GRF7 | GRF8 |
| --- | --- | --- | --- | --- | --- | --- | --- | --- |
| **4sgRNA sample** |  |  |  |  |  |  |  |  |
| Mapping rate | .97 | .97 | .98 | .95 | .97 | .96 | .97 | .91 |
| Mapped reads | 114483 | 123969 | 99432 | 106500 | 114976 | 92153 | 84857 | 77923 |
| Reads per copy<sup>1</sup> | 50 | 54 | 43 | 46 | 50 | 40 | 37 | 34 |
| Basis<sup>2</sup> | 52662 | 57026 | 49716 | 53250 | 52889 | 42390 | 42429 | 38962 |
| Indels<sup>3</sup> | 1112 | 45027 | 1277 | 22327 | 3872 | 16865 | 0 | 5779 |
| Mutation rate<sup>4</sup> | .02 | .79 | .03 | .42 | .07 | .40 | 0 | .15 |
|  | .02 | .83 | .03 |  | .07 | .47 | 0 | .12 |
| **8sgRNA sample** |  |  |  |  |  |  |  |  |
| Mapping rate | 0.95 | 0.94 | 0.97 | 0.93 | 0.97 | 0.94 | 0.97 | 0.96 |
| Mapped reads | 79337 | 120604 | 145741 | 128946 | 134287 | 59646 | 208216 | 133091 |
| Reads per copy<sup>1</sup> | 29 | 46 | 56 | 49 | 51 | 23 | 79 | 51 |
| Basis<sup>2</sup> | 76957 | 116986 | 141369 | 125078 | 130258 | 57857 | 201970 | 129098 |
| Indels<sup>3</sup> | 871 | 89789 | 330 | 35892 | 5801 | 18986 | 0 | 14059 |
| Mutation rate<sup>4</sup> | .01 | .77 | <.01 | .29 | .04 | .32 | 0 | .11 |
|  | .01 | .85 | .01 |  | .05 | .45 | 0 | .16 |

<sup>1</sup> The number of reads expected to be generated by one allele of one seedling in the sample; calculated by dividing *mapped reads* by two-times the number of seedlings in the sample (2 x 1151 for '4sgRNA'; 2 x 1310 for '8sgRNA').
- <sup>2</sup> The number of reads expected from seedlings that had been mutagenized for a given GRF locus. In the '4sgRNA' sample, 527 seedlings (46%) were from parents harboring the '1256' construct, 582 (50%) from parents harboring the '3478' construct, and 42 (4%) were control seedlings; *basis* is calculated as *mapped reads* × 0.46 for GRF1, GRF2, GRF5, GRF6, and *mapped reads* × 0.5 for GRF3, GRF4, GRF7, GRF8. In the '8sgRNA' sample, 1265 seedlings (97%) were from parents harboring both constructs, and 45 (3%) were control seedlings; thus, *basis* is calculated as *mapped reads* × 0.97. - <sup>3</sup> Insertion-deletion events detected by AGEseq; events supported by fewer reads than *reads per allele* were considered artifacts and not included. - <sup>4</sup> Calculated as *indels* / *basis*. To validate the estimates, mutation frequencies were also estimated using the mapping and SNP-calling functions of Geneious 10.1.2 (https://www.geneious.com); the results are listed in cursive below the AGEseq estimates.

## Supporting information

Annotated listings of plasmid sequences

Description of LED illuminations

List of oligonucleotides

## Supporting Information

**S1 File. Annotated sequence listing of T-DNA vectors**. (zip format)

**S2 File. LED illumination**. (pdf format)

**S3 File. Oligonucleotides for plasmid construction and PCR**. (pdf format)

## Acknowledgements

We thank Lisa Donovan (U of Georgia, USA) for her generous support of the PBIO3660L lab course; David Nelson (U of California Riverside, USA) for bringing the pHEE401E toolbox to our attention; Chengchao Zheng (Shandong Agricultural U, China) and Bob Schmitz (U of Georgia, USA) for kindly sharing pHEE401E and pCBC-DT1DT2 DNA; Magdy Alabady (U of Georgia, USA) for advice on designing primers for amplicon sequencing; Liangjiao Xue (Nangjing Forestry U, China) for help with installing the AGE-seq package; Chad Niederhuth (Michigan State U, USA) and Cordula Schulz (U of Georgia, USA) for valuable comments on the manuscript.

## Author Contributions

*Conceptualization*: Wolfgang Lukowitz

*Data Curation*: n/a

*Formal Analysis*: n/a

*Funding Acquisition*: n/a

*Investigation*: Juan Angulo, Christopher P Astin, Olivia Bauer, Kelan J Blash, Natalee M Bowen, Nneoma J Chukwudinma, Austin S Dinofrio, Donald O Faletti, Alexa M Ghulam, Cloe M Gusinde-Duffy, Kamaria J Horace, Andrew M Ingram, Kylie E Isaack, Geon Jeong, Randolph J Kiser, Jason S Kobylanski, Madeline R Long, Wolfgang Lukowitz, Grace A Manning, Julie M Morales, Kevin H Nguyen, Robin T Pham, Monthip H Phillips, Tanner W Reel, Jenny E Seo, Hiep D Vo, Alexander M Wukuson, Kathryn A Yeary, Grace Y Zhang

*Methodology*: n/a

*Project Administration*: Wolfgang Lukowitz

*Resources*: n/a

*Software*: n/a

*Supervision*: Wolfgang Lukowitz

*Validation*: Wolfgang Lukowitz

*Visualization*: Wolfgang Lukowitz

*Writing – Original Draft Preparation*: Wolfgang Lukowitz

*Writing – Review & Editing*: Juan Angulo, Christopher P Astin, Olivia Bauer, Kelan J Blash, Natalee M Bowen, Nneoma J Chukwudinma, Austin S Dinofrio, Donald O Faletti, Alexa M Ghulam, Cloe M Gusinde-Duffy, Kamaria J Horace, Andrew M Ingram, Kylie E Isaack, Geon Jeong, Randolph J Kiser, Jason S Kobylanski, Madeline R Long, Wolfgang Lukowitz, Grace A Manning, Julie M Morales, Kevin H Nguyen, Robin T Pham, Monthip H Phillips, Tanner W Reel, Jenny E Seo, Hiep D Vo, Alexander M Wukuson, Kathryn A Yeary, Grace Y Zhang

## Notes

### Competing Interest Statement

The authors have declared no competing interest.

