## Supplementary material for "Targeted mutagenesis of the *Arabidopsis* GROWTH-REGULATING FACTOR (GRF) gene family suggests competition of multiplexed sgRNAs for Cas9 apoprotein": Description of LED illuminations

### Low-cost LED illumination for imaging bio-fluorescence with dissecting scopes

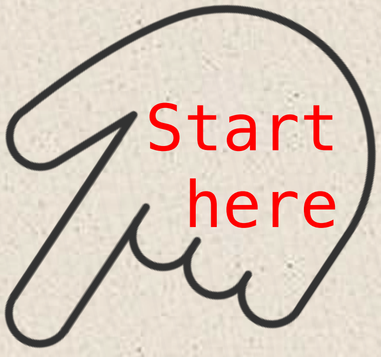


LED quad module

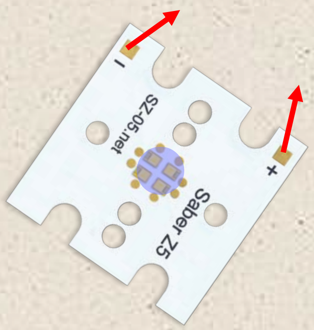


4 LED lights (●) are assembled on a 20 mm square aluminum base. Pads marked with red arrows ( ➚ ) connect to power.

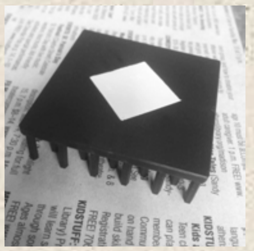

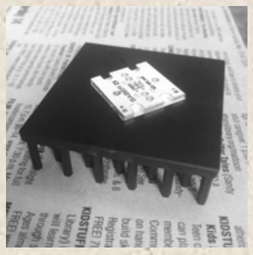

The Quad module is fixed to a 50 mm square heat sink using pre-cut thermal tape.

Soldering dots connect the power cables.

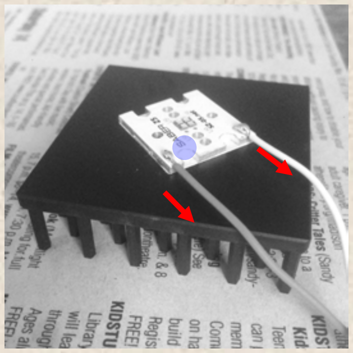

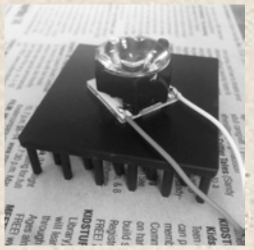

A plastic beam optics fixed with small pieces of thermal tape helps focusing emitted light.

Plugs from an extension cord for AC-adapters make swapping out lamps easy.

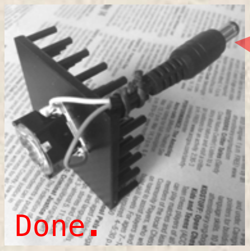


### Parts list

**LED assemblies** – Luxeon Star, www.luxeonstar.com:

- for imaging CFP: ‘violet’, SZ-05-U9, $74.28
- for imaging YFP: ‘royal blue’, SZ-05-H4, $12.20
- for imaging ‘Tomato’: ‘cyan’, SZ-05-H2, $13.00

Heat sink ($10^{\circ}$C/W or better), Luxeon Star N50-15B, $6.68
Thermal adhesive pads, set of 12, Luxeon Star LXT-SZ04-12, $7.49
Carclo 20 mm optic holder (black), Luxeon Star 10431, $0.50
Carclo $8.7^{\circ}$ circular beam optic, Luxeon Star 10193, $2.60
LED driver – Digi-Key, digikey.com, RACD06-500-LP, $10.00

**Colored-glass alternative filters** – Newport, www.newport.com:

- for imaging CFP: 50 mm square, 20CGA-475, $94.00
- for imaging YFP: 50 mm square, 20CGA-530, $94.00
- for imaging ‘Tomato’: 50 mm square, 20CGA-610, $94.00

AC-adapter extension cable, SuperBrightLEDs.com, CPS-EXT1, $1.49
Cord set with switch, Amazon, $4.00
Flexible arm cell phone holder with clamp, Walmart, $7.00

Prices as listed online July 2020.
Tools: wire cutter or knife, screw driver, soldering iron

Lamps can be mounted in the clamp of a cell phone holder.

Square emission filters can be fixed to the scope with tape.

Alternatively, 25 mm round filters can be placed on the eyepieces.

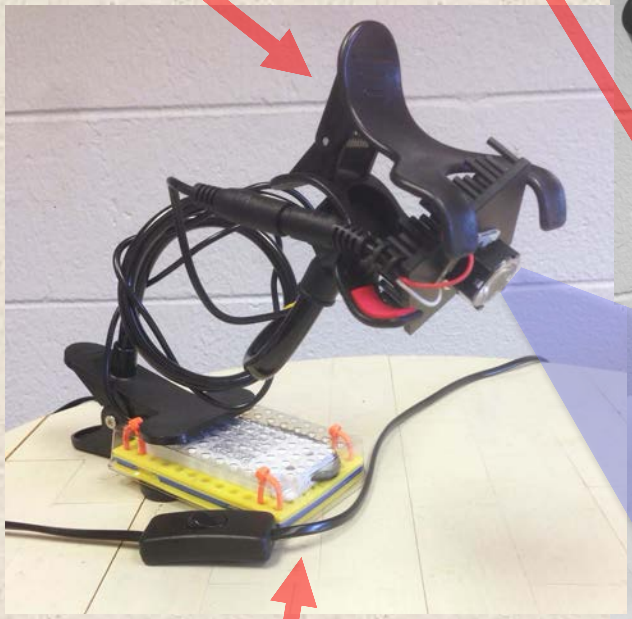

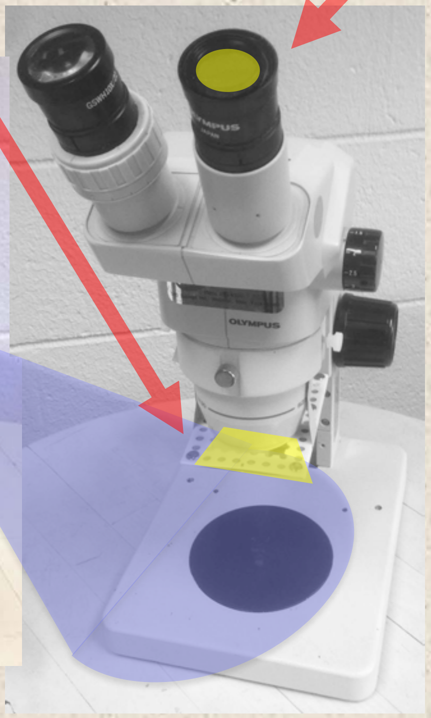


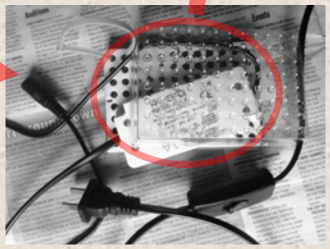

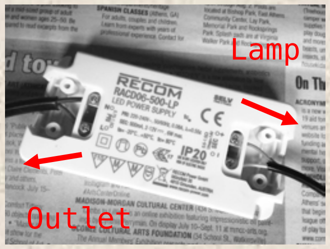


An LED driver providing 500 mA constant current (and hidden by some random plastic) powers the lamp.

For more information on performance and technical details please look up Angulo & al., CRISPR/Cas9 mutagenesis of the Arabidopsis GRF family, 2020.
