## Supplementary material for "Targeted mutagenesis of the *Arabidopsis* GROWTH-REGULATING FACTOR (GRF) gene family suggests competition of multiplexed sgRNAs for Cas9 apoprotein": List of oligonucleotides

### S3 file: Oligonucleotides for plasmid construction and PCR

#### Content

Oligonucleotides encoding the spacer sequence of T-DNAs expressing one sgRNA ... page 2

PCR-based markers for detecting CRISPR/Cas9-induced mutations ... page 3

PCR products for constructing T-DNAs expressing two sgRNAs ... page 4

PCR products for constructing T-DNAs expressing four sgRNAs ... page 5

PCR primers for mass-sequencing GRF amplicons ... page 6

**Oligonucleotides encoding the spacer sequence of T-DNAs expressing one sgRNA**

The restriction sites targeted by the sgRNAs are listed next to each GRF gene and highlighted in cyan. The nine pairs of oligonucleotides anneal to form short, double-stranded fragments with 5'-overhanging ends (5'-attg and 5'-tttg) that are compatible with the two ends generated by BsaI digestion of the T-DNA vectors. The 20 nucleotides encoding the spacer portion of the sgRNAs are listed in upper case letters. Transcription of the sgRNA invariably starts on the "g" highlighted in blue; in case of GRF2, GRF4, GRF8 and GRF9, this nucleotide forms part of the spacer sequence, in the other cases is does not match the target.

GRF1 CTGCAG (XhoI)
attgAAAGAAATGGCGGTGCTCGA n=24
aaacTCGAGCACCGCCATTTCTTT

GRF2 TCATGA (BspHI)
attGGAGAAGCAGATCTCATGAG n=23
aaacCTCATGAGATCTGCTTCTC

GRF3 CTGCAG (PstI)
attgTTTGGTATCTTAGCTGCAGA n=24
aaacTCTGCAGCTAAGATACCAAA

GRF4 CAGCTG (PvuII)
attGATGGGAAACTTCTTCAGCT n=23
aaacAGCTGAAGAAGTTTCCCAT

GRF5 GAGCTC (SacI)
attgCTAATGGAGAAGATGAGCTC n=24
aaacGAGCTCATCTTCTCCATTAG

GRF6 AGCT (AluI)
attgAGTACTTGAACACAAGAGCT n=24
aaacAGCTCTTGTGTTCAAGTACT

GRF7 CAATTG (MunI)
attgTTTCACAAACGCACAATTGA n=24
aaacTCAATTGTGCGTTTGTGAAA

GRF8 CAGCTG (PvuII)
attGACAGGAGGCTCATTGCAGC n=23
aaacGCTGCAATGAGCCTCCTGT

GRF9 GGCC (HaeIII)
attGGATGAGGAATAAGTGGCCG n=23
aaacCGGCCACTTATTCCTCATC

### PCR-based markers for detecting CRISPR/Cas9-induced GRF mutations

The approximate size of the PCR fragments as well as the restriction products are listed for each GRF gene. Induced mutations are predicted to eliminate the restriction sites. For lack of alternatives, the GRF6 marker targets an AluI site (agct). A "g" of the reverse primer (listed in red and highlighted in yellow) is mismatched to the genomic target sequence (harboring an "a" at the corresponding position) in order to eliminate a neighboring AluI site; the diagnostic bands of this marker are relatively small and were resolved on a 3% gel prepared with a high-resolution blend agarose (Amresco 3:1 HRB, E776). Germline transmitted GRF alleles were sequenced using either one of the PCR primers or an internal sequencing primer.

GRF1 ~750 bp; XhoI-digested: ~500 + 250 bp
GCATGGAAACTTGTTGACAGG n=21
GTTGTGTTCCAACAGCAGC n=19

GRF2 ~610 bp; BspHI-digested: ~480 + 130 bp
GGTGAGGATTGCTTGCAACG n=20
GGAGGCAAAGATCCGTAAGG n=20
Internal sequencing primer: AGGAATCTGGTGAAGAAACGG n=21

GRF3 ~480 bp; PstI-digested: ~410 + 70 bp
GCAACTGAAACAATGGAGAAGC n=22
GGTAGGTGTTGAAGAGGATGG n=20
Internal sequencing primer: TGCCCAGCTAAAGAAGC n=17

GRF4 ~890 bp; PvuII-digested: ~650 + 240 bp
CTGCTACTTCTGCTGCTGC n=19
GTTGTGATGGTGGTTGTGG n=19

GRF5 ~730 bp; SacI-disgested: ~600 + 130 bp
TGAGTCTAAGTGGAAGTAGCG n=21
GAAGACAAGGTGGGACTTGG n=20

GRF6 ~230 bp; AluI-digested: ~210 + 20 bp
CATGAAGCACAGAGATTCTGC n=21
GCATATTTGCgGCTAAGTACTTGAAC n=26

GRF7 ~560 bp; MunI-digested: ~390 + 170 bp
GAGACATGGAGATTCATCCGC n=21
GATGATGAAACCTCCATTGGC n=21

GRF8 ~580 bp; PvuII-digested: ~300 + 280 bp
CACCCTTTCCCAAGAACCAC n=20
GATCTGGCTTATAGGTTGTCCG n=22

GRF9 ~650 bp; HaeIII-digested: 420 + 230 bp
TGAACAGTAGCGAGCAGAGC n=20
TCCAACTCAGAACCCAAAAGC n=21

**PCR products for constructing T-DNAs expressing two sgRNAs**

All PCR fragments were produced by pairs of nested primers with a proofreading enzyme (Q5 polymerase, New England Biolabs, M0491). Inner primers (50 nM f.c.) annealed to the template plasmids and in addition encoded the spacer portion of the sgRNAs. Outer primers (1 µM f.c., 20-fold excess of inner primers) used the DNA fragments produced by the inner primers during the first few PCR cycles as template and in addition contained sequences required for generating vector-compatible ends by BsaI digestion. The BslI recognition sequence (ggtcttn$_{1/5}$) is highlighted in green, the start sites of small RNA transcripts in blue, and the nucleotides corresponding to the genomic restriction sites targeted by the CRISPR/Cas9 complexes in cyan. Primers marked by grey highlights were also used for constructing T-DNAs expressing four sgRNAs.

Two plasmids were used as PCR templates: pGEM-2t for constructs harboring a single polycistronic small RNA gene (with the two sgRNAs separated by an alanine tRNA); and pCBC-DT1T2 for constructs harboring two separate sgRNA genes.

PCR fragments for single small RNA gene with tRNA spacer
~200 bp; produced by GRF_-iF/-iRT (50 nM) and GRF_-2F/-2R from pGEM-2T

PCR fragments for tandem array of two sgRNA genes
~600 bp; produced by GRF_-iF/-iRT (50 nM) and GRF_-2F/-2R from pCBC-DT1DT2

| | | |
|---|---|---|
| GRF1-2F | atatatggtctcgattgAAAGAAATGGCGGTGCTCGAgtt | n=40 |
| GRF1-iF | AAAGAAATGGCGGTGCTCGAgttttagagctagaaatagc | n=40 |
| GRF2-iRP | aacCTCATGAGATCTGCTTCTCCaatctcttagtcgactctac | n=43 |
| GRF2-iRT | aacCTCATGAGATCTGCTTCTCCtgcaccagccgggaatc | n=40 |
| GRF2-2R | attattggtctcgaaacCTCATGAGATCTGCTTCTCC | n=37 |
| GRF3-2F | atatatggtctcgattgTTTGGTATCTTAGCTGCAGAgtt | n=40 |
| GRF3-iF | TTTGGTATCTTAGCTGCAGAgttttagagctagaaatagc | n=40 |
| GRF4-iRP | aacAGCTGAAGAAGTTTCCCATCaatctcttagtcgactctac | n=43 |
| GRF4-iRT | aacAGCTGAAGAAGTTTCCCATCtgcaccagccgggaatc | n=40 |
| GRF4-2R | attattggtctcgaaacAGCTGAAGAAGTTTCCCATC | n=37 |
| GRF5-2F | atatatggtctcgattgCTAATGGAGAAGATGAGCTCgtt | n=40 |
| GRF5-iF | gCTAATGGAGAAGATGAGCTCgttttagagctagaaatagc | n=41 |
| GRF6-iRP | aacAGCTCTTGTGTTCAAGTACTCaatctcttagtcgactctac | n=44 |
| GRF6-iRT | aacAGCTCTTGTGTTCAAGTACTCtgcaccagccgggaatc | n=41 |
| GRF6-2R | attattggtctcgaaacAGCTCTTGTGTTCAAGTACTC | n=38 |
| GRF7-2F | atatatggtctcgattgTTTCACAAACGCACAATTGAgtt | n=40 |
| GRF7-iF | gTTTCACAAACGCACAATTGAgttttagagctagaaatagc | n=41 |
| GRF8-iRP | aacGCTGCAATGAGCCTCCTGTCaatctcttagtcgactctac | n=43 |
| GRF8-iRT | aacGCTGCAATGAGCCTCCTGTCtgcaccagccgggaatc | n=40 |
| GRF8-2R | attattggtctcgaaacGCTGCAATGAGCCTCCTGTC | n=37 |

### PCR products for constructing T-DNAs expressing four sgRNAs

The pTUU-1256 and pYUU-3478 T-DNAs were assembled from the BsaI-linearized vector backbone and three PCR fragments using an NEBuilder kit (New England Biolabs, E2621). A proofreading enzyme (Q5 polymerase, New England Biolabs, M0491) was used for generating the PCR-fragments, and some reactions contained nested primers at a low concentration (50 nM, a 20$^{th}$ of the other primers) as described above. The portions of PCR primers creating the overlaps required for fragment assembly are underlined below, and the length of the overlaps is noted in brackets. Primer sequences encoding the spacer portions of the sgRNAs are listed in uppercase. Cyan highlights mark nucleotides that correspond to the genomic restriction sites targeted by the CRISPR/Cas9 complexes, and blue highlights mark the start sites of small RNA transcripts. Six of the primers were also used for constructing T-DNAs expressing two sgRNAs (grey highlights).

#### pTUU-1256

~240 bp; produced by GRF1-iF (50 nM), GRF1-4F, GRF2-iRT from pGEM-2t as template
~600 bp; produced by GRF2-4F and GRF5-4R from pCBC-DT1DT2 as template
~230 bp; produced by GRF6-iR (50 nM), GRF5-iF, GRF6-4R from pGEM-2t as template

| | | | |
|---|---|---|---|
| GRF1-4F | acagctagagtcgaagtagtgattgAAAGAAATGGCGGTGCTC | n=43 | (25 bp overlap) |
| GRF1-iF | AAAGAAATGGCGGTGCTCGAgttttagagctagaaatagc | n=40 | |
| GRF2-iRT | aacCTCATGAGATCTGCTTCTCCtgcaccagccgggaatc | n=40 | |
| | | | (23 bp overlap) |
| GRF2-4F | GGAGAAGCAGATCTCATGAGgttttagagctagaaatagc | n=40 | |
| GRF5-4R | acGAGCTCATCTTCTCCATTAGcaatctcttagtcgactc | n=40 | |
| | | | (23 bp overlap) |
| GRF5-iF | gCTAATGGAGAAGATGAGCTCgtttagagctagaaatagc | n=41 | |
| GRF6-iR | aacAGCTCTTGTGTTCAAGTACTtgcaccagccgggaatc | n=40 | |
| GRF6-4R | aacttgctatttctagctctaaaacAGCTCTTGTGTTCAAG | n=41 | (25 bp overlap) |

#### pYUU-3478

~240 bp; produced by GRF3-iF (50 nM), GRF3-4F, GRF4-iRT from pGEM-2t as template
~600 bp; produced by GRF4-4F, GRF7-4R from pCBC-DT1DT2 as template
~230 bp; GRF7-iRT (50 nM), GRF7-iF, GRF8-4R from pGEM-2t as template

| | | | |
|---|---|---|---|
| GRF3-4F | acagctagagtcgaagtagtgattgTTTGGTATCTTAGCTGCAG | n=44 | (25 bp overlap) |
| GRF3-iF | TTTGGTATCTTAGCTGCAGAgttttagagctagaaatagc | n=40 | |
| GRF4-iRT | aacAGCTGAAGAAGTTTCCCATCtgcaccagccgggaatc | n=40 | |
| | | | (23 bp overlap) |
| GRF4-4F | GATGGGAAACTTCTTCAGCTgttttagagctagaaatagc | n=40 | |
| GRF7-4R | acTCAATTGTGCGTTTGTGAAAcaatctcttagtcgactc | n=40 | |
| | | | (23 bp overlap) |
| GRF7-iF | gTTTCACAAACGCACAATTGAgttttagagctagaaatagc | n=40 | |
| GRF8-iRT | aacGCTGCAATGAGCCTCCTGTCtgcaccagccgggaatc | n=40 | |
| GRF8-4R | aacttgctatttctagctctaaaacGCTGCAATGAGCCTCC | n=41 | (25 bp overlap) |

**PCR primers for mass-sequencing GRF amplicons**

For each GRF gene, the chromosome coordinates and the length of the *Arabidopsis* genomic fragment captured in the amplicons are shown. Primers are oriented with respect to the GRF coding sequence (which is reverse complementary to the chromosome reference sequence in the case of GRF2, GRF3 and GRF4). Only the gene-specific portions of the primers are listed; all primers also contained 5'-tails for the purpose of library production:

| | | |
|---|---|---|
| Forward primer tail: | tcgtcggcagcgtcagatgtgtataagagacagn | n=33 |
| Reverse primer tail: | gtctcgtgggctcggagatgtgtataagagacag | n=34 |

GRF1 Chr2:9729795..9729995 (201 bp)
TGGGGCTCTTTTCATCTGG n=19
GGCAGCATTAGTATTGTGGC n=20

GRF2 Chr4:17727297..17727493 (264 bp)
CTTGCAACGAAGCTTGAAGC n=20
CGTCTGGTTTATCTGAGAAGC n=21

GRF3 Chr2:15272424..15272583 (160 bp)
AGCAGCAACAACATCAGAC n=19
TCAGGAGACAAAGAGGAGAG n=20

GRF4 Chr3:19617858..19618055 (198 bp)
TCTTCTTCCAGGTTCCTCAG n=20
GCACCAGCCAACATGTATC n=19

GRF5 Chr3:4608577..4608736 (160 bp)
ACACCAACACAATGGGAAG n=19
TATAGGGCTTACGGGATTGG n=20

GRF6 Chr2:2426350..2426520 (170 bp)
GGATTCCATTCACAGAATCAC n=21
GGGAGAGAAGAAGCTTGAAG n=20

GRF7 Chr5:21794856..21795043 (188 bp)
CAGGAAATGGACTTTTGGG n=19
CTCTAAATCTCCGGCGATG n=19

GRF8 Chr4:12538390..12538569(180 bp)
TAGTGGATACATATAGGAGTGG n=22
CTGTTCTCTTGCACCTCC n=18

GRF9 Chr2:18745573..18745750 (178 bp)
TGTTATGAAGATGCAGAGCC n=20
CCAAATAGGCACCACGAG n=18
